# A palmitate-rich metastatic niche enables metastasis growth via p65 acetylation

**DOI:** 10.1101/2022.10.24.513556

**Authors:** Patricia Altea-Manzano, Ginevra Doglioni, Alejandro M. Cuadros, Emma Nolan, Juan Fernandez-Garcia, Qi Wu, Florencia Cidre-Aranaz, Aurelie Montagne, Mélanie Planque, Oskar Marin-Bejar, Joke Van Elsen, Ines Vermeire, Dorien Broekaert, Carla Riera-Domingo, François Richard, Tatjana Geukens, Maxim De Schepper, Sophia Leduc, Sigrid Hatse, Yentl Lambrechts, Emily Jane Kay, Sergio Lilla, Sofie Demeyer, Vincent Geldhof, Bram Boeckx, Alisa Alekseenko, Celia de la Calle Arregui, Giuseppe Floris, Jean-Christophe Marine, Diether Lambrechts, Vicent Pelechano, Massimiliano Mazzone, Sara Zanivan, Jan Cools, Hans Wildiers, Véronique Baud, Thomas G.P. Grünewald, Christine Desmedt, Ilaria Malanchi, Sarah-Maria Fendt

**Affiliations:** Laboratory of Cellular Metabolism and Metabolic Regulation, VIB-KU Leuven Center for Cancer Biology, VIB, Herestraat 49, 3000 Leuven, Belgium; Laboratory of Cellular Metabolism and Metabolic Regulation, Department of Oncology, KU Leuven and Leuven Cancer Institute (LKI), Herestraat 49, 3000 Leuven, Belgium; The Francis Crick Institute. 1 Midland Road, London NW1 1AT, United Kingdom; Laboratory of Experimental Oncology, Department of Oncology, KU Leuven. Herestraat 49 - box 815 3000 Leuven, Belgium; Hopp-Children’s Cancer Center (KiTZ), Heidelberg, Germany; Division of Translational Pediatric Sarcoma Research, German Cancer Research Center (DKFZ), German Cancer Consortium (DKTK), Deutsches Krebsforschungszentrum (DKFZ) Div. Translational Pediatric Sarcoma Research/B410 Im Neuenheimer Feld 280, 69120 Heidelberg, Germany; Université de Paris, NF-kappaB, Différenciation et Cancer, F-75006 Paris, France; Laboratory for Molecular Cancer Biology, Center for Cancer Biology, VIB; Laboratory for Molecular Cancer Biology, Department of Oncology, KU Leuven. Herestraat 49 - box 912 3000 Leuven, Belgium; Laboratory of Tumor Inflammation and Angiogenesis (VIB-KULeuven) Herestraat 49, Box 912 ON IV, 3000 Leuven Belgium; Laboratory for Translational Breast Cancer Research, Department of Oncology, Herestraat 49 - box 815 3000 Leuven, Belgium; Cancer Research UK Beatson Institute. G61 1BDUK, Glasgow, UK; Laboratory for molecular biology of leukemia (VIB-KULeuven) Herestraat 49, Box 912 ON IV, 3000 Leuven, Belgium; Laboratory for angiogenesis and vascular metabolism (VIB-KULeuven) ON IV, 3000 Leuven Belgium; Laboratory of Translational Genetics, VIB Center for Cancer Biology, 3000 Leuven, Belgium; Laboratory of Translational Genetics, Department of Human Genetics, KU Leuven, 3000 Leuven, Belgium; SciLifeLab, Department of Microbiology, Tumor and Cell Biology. Karolinska Institute, Tomtebodavägen 23A, Box 1031, 171 65 Solna, Sweden; Department of Imaging and Pathology, Laboratory of Translational Cell & Tissue Research and University Hospitals Leuven, Department of Pathology, KU Leuven, Herestraat 49, Box 912 ON IV, 3000 Leuven Belgium; Institute of Cancer Sciences, University of Glasgow, G611QH, Glasgow, UK; Institute of Pathology, Heidelberg University Hospital, Im Neuenheimer Feld 224 69120 Heidelberg Germany

**Keywords:** metastasis formation, fatty acids, palmitate, CPT1a, acetylation, NF-κB, p65, RelA, KAT2a, GCN5, breast cancer, pre-metastatic niche, AT2, obesity

## Abstract

Cancer cells outgrowing in distant organs of metastasis rewire their metabolism to fuel on the available nutrients. While this is often considered an adaptive pressure limiting metastasis formation, some nutrients available at the metastatic site naturally or through changes in organ physiology may inherently promote metastatic growth. We find that the lung, a frequent site of metastasis, is a lipid-rich environment. Moreover, we observe that pathological conditions such as pre-metastatic niche formation and obesity further increase the availability of the fatty acid palmitate in the lung. We find that targeting palmitate processing inhibits spheroid growth *in vitro* and metastasis formation in lean and obese mice. Mechanistically, we discover that breast cancer cells use palmitate to synthesize acetyl-CoA in a carnitine palmitoyltransferase 1a (CPT1a)-dependent manner. Lysine acetyltransferase 2a (KAT2a), whose expression is promoted by palmitate availability, relies on the available acetyl-CoA to acetylate the NF-κB subunit p65. This favors nuclear location of p65 and activates a pro-metastatic transcriptional program. Accordingly, deletion of KAT2a phenocopies CPT1a silencing *in vitro* as well as *in vivo* and patients with breast cancer show co-expression of both proteins in metastases across palmitate-rich metastatic sites. In conclusion, we find that palmitate-rich environments foster metastasis growth by increasing p65 acetylation resulting in elevated NF-κB signaling.

## Introduction

Nutrient availability is a key aspect of a permissive environment enabling metastasis formation ^1,2^. Certain nutrients such as glucose ^3,4^, fatty acids ^5,6^, pyruvate ^7-9^ and glutamine ^10^ are aiding metastasizing cancer cells when they are seeding and colonizing in a distant organ. While nutrient availability is certainly defined by the functional processes occurring in healthy organs, the question arises whether aberrant disease processes or dietary conditions can influence the nutrient spectrum or concentrations available in organs of metastasis.

A nutrient class highly linked to metastasis formation are fatty acids, which are also available through diet ^11-13^. Accordingly, feeding mice with a high-fat diet increases metastasis formation in multiple cancers such as breast and oral cancers ^14-17^. In many cancer types, blocking fatty acid uptake by targeting the lipid receptor CD36 is sufficient to impair metastasis formation ^11^. Moreover, fatty acids found in high fat diets such as palmitate and oleate promote cancer progression of oral ^16,17^ and melanoma ^18^ tumors, respectively. However, it remains largely elusive whether these fatty acids are available in organs of future metastasis and whether obesity resulting from high fat diet feeding alters their concentration.

Another aberrant disease process directly linked to cancer progression is pre-metastatic niche formation ^19^. In this process, tumor secreted factors prime the environment of a future organ of metastasis to create a permissive environment. In particular, there is extensive evidence that the immune cell and extracellular matrix composition of the pre-metastatic niche are highly shaped by tumor secreted factors resulting in the increased success of cancer cells in seeding and colonizing in such primed organs ^19,20^. However, to date very little is known about nutrient priming of the pre-metastatic niche with only one report showing increased glucose availability resulting from the hypometabolism of resident lung cells induced by the tumor secreted microRNA miR-122 ^21^.

Here we analyzed the fatty acid composition of healthy lungs and found that it is a lipid-rich environment. In addition, we observed that mice exposed to high fat diet or undergoing pre-metastatic niche formation show a significant increase in lung palmitate availability. We further show that breast cancer cells seeding and colonizing the lung rely on CPT1a for fatty acid oxidation, which in turn activates NF-κB signaling via the acetylation of p65 in a KAT2a-dependent manner. Accordingly, CPT1a and KAT2a targeting is highly effective in inhibiting metastasis formation *in vivo*.

## Results

### The healthy lung is a lipid-rich environment and palmitate availability further increases upon obesity and pre-metastatic niche formation

Although the lung is a frequent metastatic site, its lipid concentration remains unknown. We isolated lung interstitial fluid, which is a local source of nutrients for metastasized cancer cells, from healthy Balb/c mice and non-cancerous lung tissue of human individuals ^7,22^. Subsequently, we measured fatty acid availability using mass spectrometry. We found that mouse and human lung interstitial fluids were very similar regarding fatty acid composition and concentrations, and remarkably, palmitate oleate and linoleate were highly available (**Figure 1A**).

**Figure 1.**
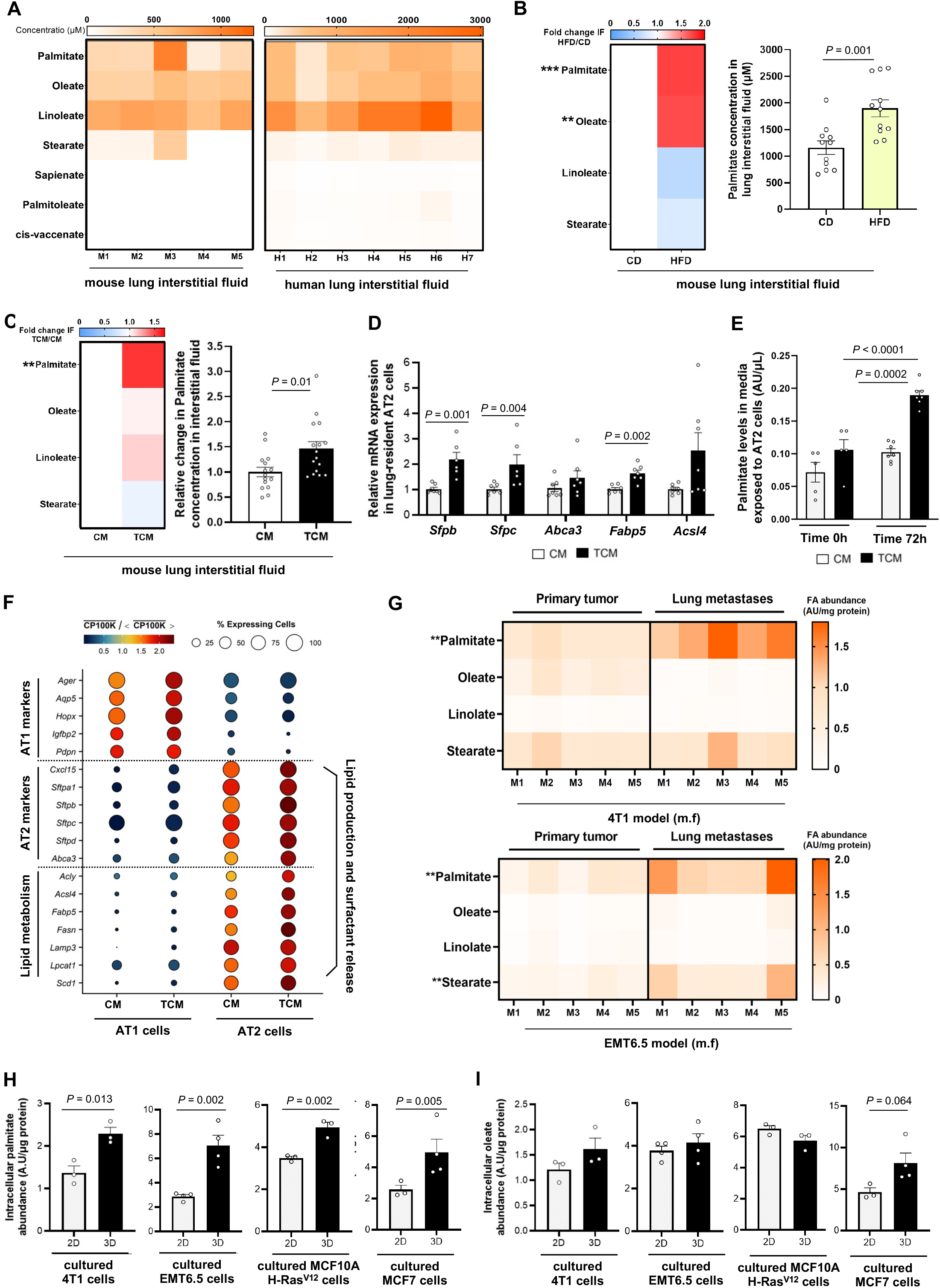
Palmitate availability in the lung is increased upon high fat diet and pre-metastatic niche formation. **A**. Fatty acid concentrations in the lung interstitial fluid of healthy BALB/c mice (n = 5) and human patients (n = 7). **B**. Relatives changes in fatty acid concentrations in lung interstitial fluid of BALB/c mice after 16 weeks on control diet (CD, 13%/27%/60% fat/protein/carbohydrates) or high fat diet (HFD, 60%/20%/20% fat/protein/carbohydrates). Left panel represents fold changes compared concentration found in lung of mice fed with high fat diet (HFD) and control diet (n = 11). e Right panel represents palmitate concentration. Data are presented as absolute concentration measured by mass spectrometry. Two-tailed unpaired Student’s t test (n =11). Asterisks represent statistical significance (**p ≤ 0.01 and ***p ≤ 0.001) **C**. Relatives changes in fatty acid concentrations in lung interstitial fluid of healthy BALB/c mice after periodic intravenous injections of control media (CM) or tumor conditioned media (TCM). The right panel represents palmitate concentration as fold change compared with control media (CM). Two-tailed unpaired Student’s t test. n = 16. Asterisks represent statistical significance (**p ≤ 0.01) **D**. Relative expression of genes involved in pulmonary surfactant production and secretion in alveolar type II (AT2) cells exposed to control (CM) or tumor conditioned media (TCM) for 72h. **E**. Palmitate levels present in lung resident AT2 cells exposed for 0 and 72h to control (CM) or tumor conditioned (TCM) media. Two-tailed unpaired student’s T-test (n = 6). **F**. Gene expression vs. cell type and condition media profiles for known marker for AT1, AT2 cells and lipid-related genes indicated on the left-hand side. Scaled expression levels are indicated by the color scale, where (CP100k) □denotes the average gene expression level (in counts per 100k reads) over all cells of a given type in each condition, and ⟨(CP100k) □⟩ the average of the latter over all cell types and condition media. The areas of the circles represent the percentage of cells with non-zero expression of each gene among all cells of each type and in each condition media. **G**. Fatty acid abundance in 4T1 and EMT6.5 primary tumor tissues and the matching lung metastases. Data represent the metabolite ion counts normalized by the amount of protein. Two-sided Mann-Whitney test (n = 5). Asterisks represent statistical significance (**p ≤ 0.01). **H** and **I.** Intracellular palmitate and oleate abundance from mouse (4T1, EMT6.5) and human (MCF10A H-Ras^V12^, MCF7) breast cancer cells cultured on soft-agar (3D) or attached (2D) conditions. Two-tailed unpaired student’s T-test (n ≥3).

Next, we asked whether the availability of these fatty acids changes in pathological conditions known to promote metastasis such as obesity ^15,23^ and pre-metastatic niche formation ^20^. To investigate that, we fed BALB/c mice a high fat diet (60%/20%/20% fat/protein/carbohydrates) or a control diet (13%/27%/60% fat/protein/carbohydrates) for 16 weeks (Supplemental Figure 1A). We found that lung interstitial fluid of obese mice had increased fatty acids levels ―specifically palmitate and oleate ― compared to lean mice **(Figure 1B)**. Subsequently, we investigated whether tumor secreted factors also influence fatty acid availability in the pre-metastatic niche. To mimic pre-metastatic niche formation, we generated tumor-conditioned media (TCM) by culturing mouse 4T1-derived primary breast tumors for 3 days in DMEM-F12 media ^24-27^ and injected this TCM or control media (CM) for three weeks intravenously to healthy BALB/c mice ^28,29^ **(Supplemental Figure 1B)**. TCM compared to CM injections led to a robust generation of an experimental pre-metastatic niche in the lungs as defined by changes in glucose availability, immune cells recruitment, and gene expression patterns of stromal cells as described previously ^30-35^ **(Supplemental Figure 1C-E)**. Consecutively, we isolated lung interstitial fluid from mice treated with TCM or CM and measured fatty acid concentrations using mass spectrometry. We discovered that upon experimental pre-metastatic niche formation palmitate, but not oleate or linoleate, abundance increased in lung interstitial fluid (Figure 1C). Taken together, these data suggest that obesity and pre-metastatic niche formation converge into increased lung palmitate availability.

Lung resident alveolar type II cells release palmitate during pre-metastatic niche formation

Since the response to TCM was specific to palmitate, we hypothesized that some lung resident cells respond to tumor secreted factors by releasing palmitate to the pre-metastatic niche. Alveolar type II epithelial (AT2) cells are known to naturally release lipids, which form 90% of the pulmonary surfactant in healthy lungs ^36^. Thus, we investigated whether AT2 cells respond to tumor-conditioned media by increasing the expression of lipid production and pulmonary surfactant release genes and by elevating palmitate secretion. We isolated AT2 cells from *Sftpc-CreERT2;R26R-YFP* mice, cultured them on a 3D-scaffold system ^37^ and treated them with CM or TCM generated from 4T1 primary tumors. Strikingly, we observed that AT2 cells increased the expression of *Abca3, Fabp5, Sfpb, and Sfpc*, which are genes related to lipid production and pulmonary surfactant release ^36^, after tumor-conditioned media exposure **(Figure 1D)**. Moreover, we observed an elevated secretion of palmitate, but not oleate, from AT2 cells treated with TCM compared to CM **(Figure 1E, Supplemental Figure 1F)**. Hence, this shows that AT2 cells respond *in vitro* to tumor secreted factors by releasing more palmitate.

To verify this finding *in vivo*, we performed single-cell RNA sequencing (scRNA-seq) using single-cell suspensions of whole lungs generated from 10 weeks-old BALB/c mice that were injected with TCM or CM as described above. We identified the AT2 cell population based on its higher expression of the molecular markers associated with an AT2 phenotype such as *Sftpc, Sftpb* and *Abca3* ^38^ **(Figure 1F, Supplemental Figure 1G)**. In line with our *in vitro* data, we observed that the expression of lipid production and pulmonary surfactant release genes was upregulated in the AT2 cell population in mice exposed to tumor secreted factors **(Figure 1F)**. Notably, the expression of the same genes remained unchanged in alveolar type I epithelial (AT1) cells **(Figure 1F)**. Thus, we concluded that lung resident AT2 cells respond *in vitro* and *in vivo* to secreted factors from breast tumors by increasing the expression of lipid production and pulmonary surfactant release genes, which is associated with increased palmitate release *in vitro*.

### Palmitate is enriched in lung metastases compared to primary breast tumors and promotes spheroid growth

Our data show that breast cancer cells growing in the lung have access to palmitate. A consequence of this could be that metastases are enriched in palmitate. Thus, we measured the enrichment of fatty acids in lung metastases compared to primary tumors of 4T1 or EMT6.5 breast cancer cells (m.f.p. injection; 22 days) using mass spectrometry. We found that palmitate, but not oleate or linoleate, abundance increased in lung metastases compared to primary tumor tissue in both mouse models **(Figure 1G)**. Thus, these data suggest that breast cancer-derived lung metastases may take up palmitate from the lung environment.

To determine the importance of palmitate for metastasis formation, we sought to model the observed metabolic phenotype of metastases and primary tumors using an *in vitro* system. Previously, we have exploited breast cancer cell cultures in 2D monolayer and 3D spheroids for this purpose ^8,9^. In line, we found that, like metastases, 3D cultured breast cancer cells showed palmitate, but not oleate, enrichment compared to the same breast cancer cells cultured in 2D **(Figure 1H and I)**. Subsequently, we added extra palmitate and, as a negative control, oleate to 3D cultured breast cancer cells. Specifically, 75 μM conjugated palmitate or 116 μM conjugated oleate were added on top of 10% FBS resulting in a similar final concentration of 130 μM for both fatty acids. Strikingly, the addition of extra palmitate, but not oleate, stimulated 3D spheroid but not 2D monolayer growth of 4T1 cells **(Figure 2A, Supplemental Figure 2A and B)**. Since it has been often shown that oleate can buffer the effects of palmitate in cancer cells ^39-41^, we next added the combination of both fatty acids. Unexpectedly, the growth increase observed in the presence of extra palmitate was also present when palmitate supplementation was combined with oleate addition **(Supplemental Figure 2C)**. Notably, this effect was not recapitulated by supplementing stearate (another saturated fatty acid) instead of palmitate **(Supplemental Figure 2D)**, suggesting that only palmitate promotes 3D spheroid growth of 4T1 breast cancer cells. Taken together, these data show that breast cancer-derived lung metastases are enriched in palmitate compared to the corresponding primary tumors and that increasing palmitate availability further promotes 3D spheroid growth.

**Figure 2.**
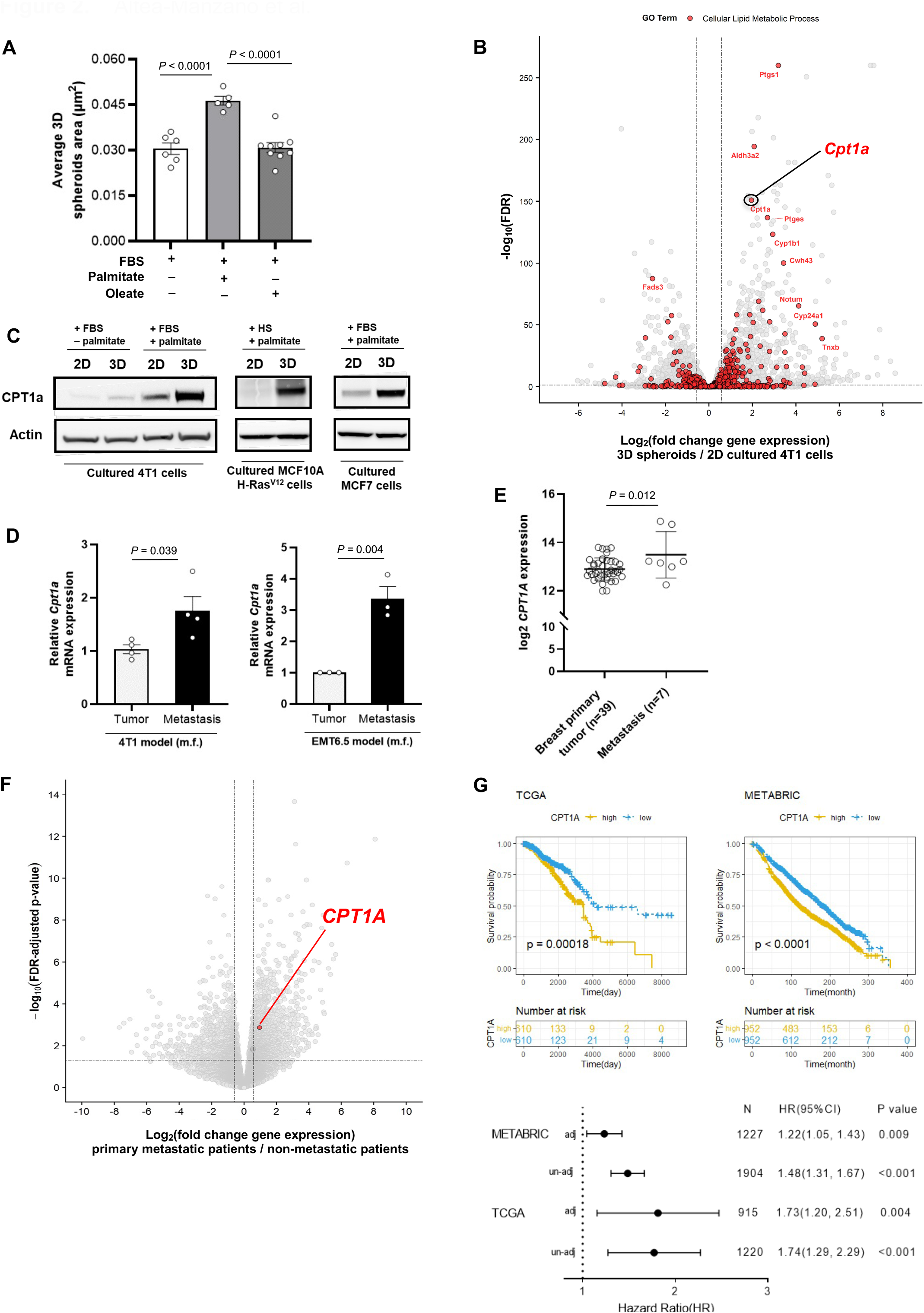
CPT1a expression is upregulated in metastasis and is associated with poor prognosis in breast cancer patients. **A**. 3D spheroids growth upon palmitate or oleate supplementation represented by the average of spheroids area of >100 spheroids (n ≥5). One-way ANOVA with Holm-Sidak’s multiple comparison test. **B**. Differentially expressed genes in 4T1 cells cultured in 3D spheroids or 2D monolayer in the presence of extra palmitate (10% FBS + palmitate). Calculated differences of gene expression are presented by plotting the negative log10 of false discovery rate (Y-axis) against the log2 fold change of gene expression (X-axis). Each dot represents an individual gene. In red, genes belong to the GEO term lipid metabolic process (GO:0006629). The highest-ranking (top 10) overexpressed genes are shown. **C**. CPT1a expression in mouse (4T1) and human (MCF10A H-Ras^V12^ and MCF7) cells growing in 2D monolayer or 3D spheroid cultures with or without additional palmitate on top of the fatty acids present in serum. A representative of three experiments is shown. **D**. Relative *Cpt1a* gene expression in 4T1 (m.f.) and EMT6.5 (m.f.) breast-derived lung metastases normalized to Cpt1a gene expression of their breast primary tumors. Two-tailed unpaired student’s T-test (n ≥3). **E**. *CPT1A* gene expression in breast primary tumors compared to metastatic tissues (GEO accession number GSE2109 (HS-00002(33)). Two-tailed unpaired Student’s t test (n ≥7). **F**. Differently expressed genes in primary tumors of primary metastatic patients (metastasis already present at diagnosis) compared to non-metastatic patients (no metastasis during at least 7 years of follow-up) from the CHEMOREL program. CPT1A transcript is identified as significantly differentially expressed (considering a FDR1<10.05) and is colored in red. **G**. Kaplan–Meier survival for breast cancer patients with high or low levels of CPT1a gene expression. Comparison of survival curves was done using Mantel-COX test and Gehan-Breslow-Wilcoxon test (n=369).

### CPT1a expression is upregulated in breast cancer-derived metastases and is associated with poor prognosis in breast cancer patients

Palmitate can have different fates in cells. Therefore, we analyzed lipid processing pathways using RNA sequencing comparing 4T1 3D spheroids versus 2D monolayers in the presence of additional palmitate and comparing 3D spheroids in the presence and absence of extra palmitate. We found the rate-limiting enzyme of the β-oxidation pathway, carnitine palmitoyltransferase 1a (CPT1a), within the highest-ranking lipid processing enzymes whose gene expression increased in 3D spheroids compared to 2D cultures and upon high compared to low palmitate availability in 3D cultures of 4T1 cells **(Figure 2B, Supplemental Figure 2E)**. We confirmed this increase at the protein level in 4T1 as well as human MCF7 and MCF10A H-Ras^V12^ breast cancer cells **(Figure 2C)**, and in lung metastases compared to the corresponding primary 4T1 and EMT6.5 breast tumors **(Figure 2D)**. Additionally, we compared our findings with the publicly available dataset Expression Project for Oncology (expO, GSE2109) and we found that CPT1a expression was higher in some metastatic samples compared to primary tumor tissue of patients with breast cancer **(Figure 2E)**.

To interrogate whether CPT1a expression is associated with progression in patients with breast cancer, we performed RNA sequencing on breast primary tumors of 14 patients that had metastases at diagnosis compared to 44 patients that did not have metastasis at diagnosis and did not develop such within at least 6-years (CHEMOREL study at UZ Leuven, see **Supplemental Table 1** for clinicopathological information). We observed that patients with metastatic breast cancer at diagnosis had higher *CPT1A* expression in their primary breast tumors compared to patients with no metastases at diagnosis **(Figure 2F)**. We further analyzed publicly available gene expression data from large cohorts of patients with breast cancer in two independent study datasets (The Cancer Genome Atlas program (TCGA) ^42^ and METABRIC ^43^). We found that high CPT1a expression (above median) in primary breast tumors of patients was significantly associated with a decreased overall survival in the TCGA and METABRIC cohort **(Figure 2G)**. This association was confirmed after correcting for the common clinic-pathological variables (age, tumor stage and tumor subtype) as detailed in **Supplemental Table 2** (HR_adj_ [95%confidence interval]: 1.73[1.20-2.51] and HR_adj_: 1.22[1.05-1.43] in TCGA and METABRIC dataset respectively). Based on these data, we concluded that CPT1a may be important for breast cancer progression in patients and that breast cancer-derived metastases may rely on CPT1a to process palmitate.

### CPT1a activity sustains palmitate-promoted spheroid growth and metastasis formation in lean and obese mice

We then assessed the effect of CPT1a inhibition in spheroid growth of breast cancer cell lines upon supplementation of extra palmitate. We found that genetic or pharmacologic inhibition of CPT1a decreased spheroid growth to the level observed without additional palmitate **(Figure 3A, 3B, Supplemental Figure 2F and G)**. Based on this *in vitro* finding, we hypothesized that blocking CPT1a, and hence intracellular palmitate utilization in the mitochondria, is sufficient to reduce lung metastases growth. To address this hypothesis, we injected 4T1 and EMT6.5 control and CPT1a knockout/down to the mammary fat pad of mice and assessed lung metastases number, area and metastatic index (metastases area divided by primary tumor weight) based on H&E staining. Strikingly, we observed that metastases number, metastases area and metastatic index were highly reduced in the absence of CPT1a expression compared to control in both mouse models **(Figure 3C-E, Supplemental Figure 3A)**, while primary tumor growth showed no changes (upon CPT1a knockdown) and a small reduction (upon CPT1a knockout) compared to control **(Supplemental Figure 3B)**. Thus, we concluded that blocking CPT1a is sufficient to impair metastatic growth in mice. Since we had seen that obesity increases palmitate levels in the lung **(Figure 1B)**, we hypothesized that CPT1a mediates the increased growth capacity of metastases in obese mice. Thus, we investigated the effect of blocking CPT1a during metastasis formation in mice with obesity **(Supplemental Figure 1A)**. We injected intravenously 4T1 and EMT6.5 control and CPT1a silenced cells expressing the congenic marker CD90.1 (as a marker for cancer cells) into lean or obese mice. Next, we measured metastatic burden by assessing the fraction of CD90.1 positive cells in the lung after 11 days. As expected, we found an increased fraction of CD90.1 positive cells in the lung of obese compared to lean mice **(Figure 3F)**. Strikingly, silencing CPT1a was sufficient to fully prevent the increase in metastatic burden induced by obesity because the fraction of CD90.1 positive cells in the lung of mice was the same comparing obese and lean mice injected with CPT1a silenced cancer cells **(Figure 3F)**. This shows that the increased growth capacity of metastases in the lung of obese mice is fully dependent on CPT1a activity.

**Figure 3.**
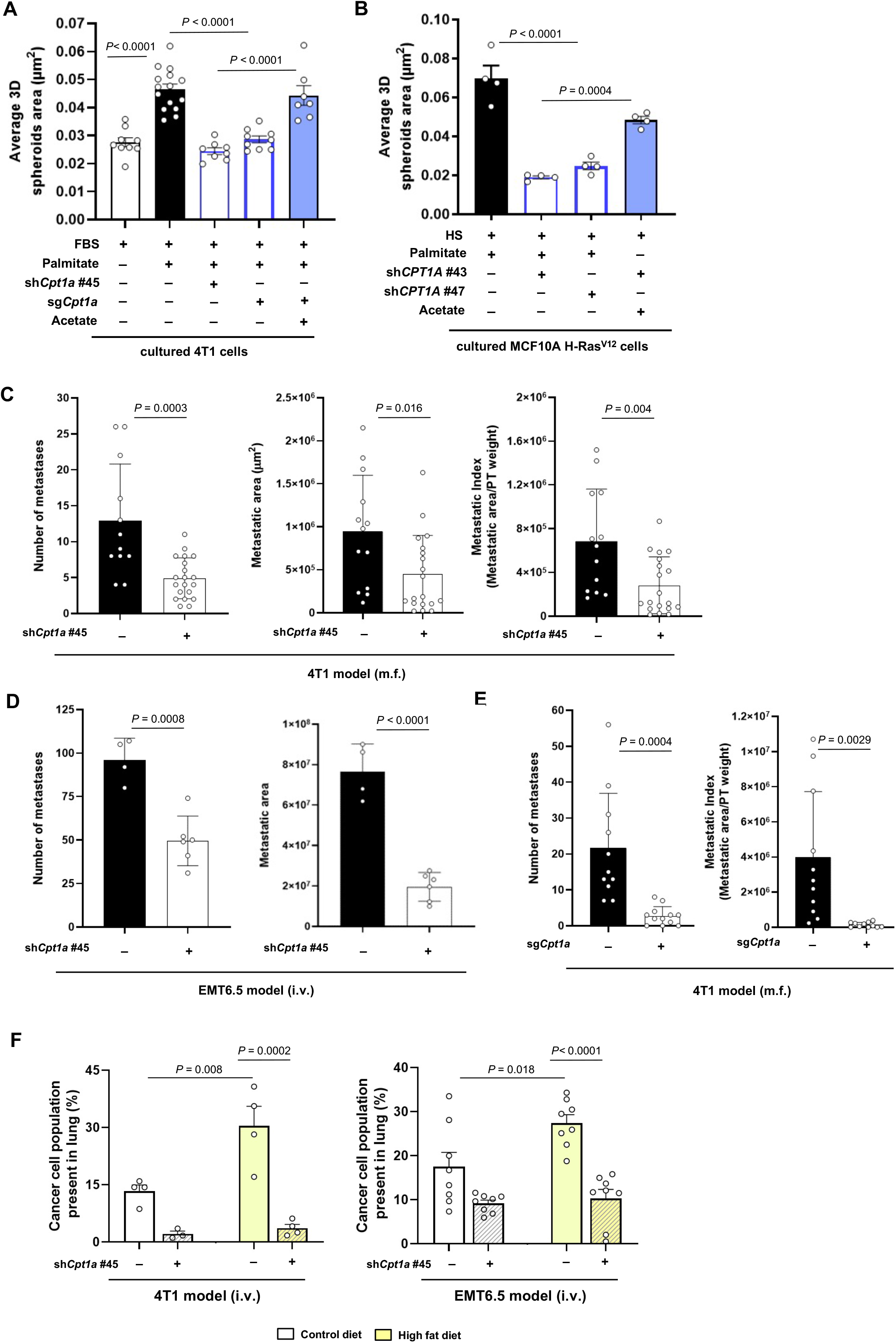
Silencing CPT1a counteracts palmitate-induced spheroid growth and inhibits metastasis formation in lean and obese mice. **A** and **B**. 3D spheroids growth of 4T1 and MCF10A H-Ras^V12^ cells upon palmitate supplementation (75 μM), CPT1a genetic inhibition performed by shRNA (shCpt1a and sh*CPT1A*) and CRISPR (sg*Cpt1a*), and upon metabolic rescue by acetate (5mM), represented by the average of spheroids area of >100 spheroids. One-way ANOVA with Turkey’s multiple comparison test (n ≥4). **C** and **D**. Metastatic burden in mice injected with 4T1 (m.f.) (**C**) and EMT6.5 (i.v.: intravenous) (**D**) cells upon genetic knockdown of Cpt1a. Two-tailed unpaired Student’s t-test (n ≥6). **E**. Metastatic burden in mice injected with 4T1 (m.f.) lungs upon genetic knockout of *Cpt1a*. m.f.: mammary fat pad. Data for control and sg*Cpt1a* are also shown in Figure 3E. Two-tailed unpaired Student’s t-test (n ≥11). **F**. Percentage of cancer cells present in lung of mice after 11 days of i.v injections with CD90.1 positive 4T1 or EMT6.5 cancer cells. Cancer cells were previously transduced with a lentiviral vector with shRNA against *Cpt1a* or scramble sequence. Before injections, mice were maintained during 16 weeks in CD and HFD. Two-way ANOVA with Turkey’s multiple comparison test (n ≥4).

### CPT1a activity sustains acetyl-CoA levels

Next, we investigated the mechanism by which CPT1a-mediated palmitate utilization supports metastasizing cancer cells. CPT1a facilitates the transport of fatty acids into the mitochondria for subsequent oxidation to acetyl-CoA. Therefore, we measured acetyl-CoA abundance in 3D spheroids and metastases tissue using mass spectrometry. We found that acetyl-CoA abundance was higher in breast cancer cells supplemented with extra palmitate in 3D compared to 2D culture **(Figure 4A and Supplemental Figure 4A)** and in lung metastases compared to primary 4T1 and EMT6.5 breast tumors **(Figure 4B and C)**. In case CPT1a activity and consecutive fatty acid oxidation are responsible for this elevated acetyl-CoA abundance in spheroids and metastases, blocking CPT1a activity would decrease acetyl-CoA abundance. To investigate this possibility, we silenced CPT1a and used the irreversible CPT1a inhibitor etomoxir in cultured breast cancer cells (50 μM) as well as in mice harboring 4T1 lung metastases (40 mg/kg, i.p. injection once a day starting 72 hours before euthanasia ^44^). In line with our hypothesis, we observed that acetyl-CoA abundance decreased in 3D cultured mouse (4T1 and EMT6.5) and human (MCF7 and MCF10A H-Ras^V12^) breast cancer cells upon CPT1a silencing **(Figure 4D and Supplemental Figure 4B)** as well as in 4T1 lung metastases when the mice were treated with etomoxir **(Figure 4C)**. In the absence of extra palmitate, CPT1a silencing did not decrease acetyl-CoA abundance in 3D cultured 4T1 breast cancer cells **(Supplemental Figure 4C)**. Moreover, the 3D growth defect observed upon CPT1a inhibition was rescued by the supplementation of acetate (5mM; **Figure 3A and B)** and octanoate (130 μM, **Supplemental Figure 4D)**, which are CPT1a independent sources of acetyl-CoA. These findings show that breast cancer spheroids in the presence of additional palmitate and breast cancer-derived lung metastases rely on CPT1a for acetyl-CoA production.

**Figure 4.**
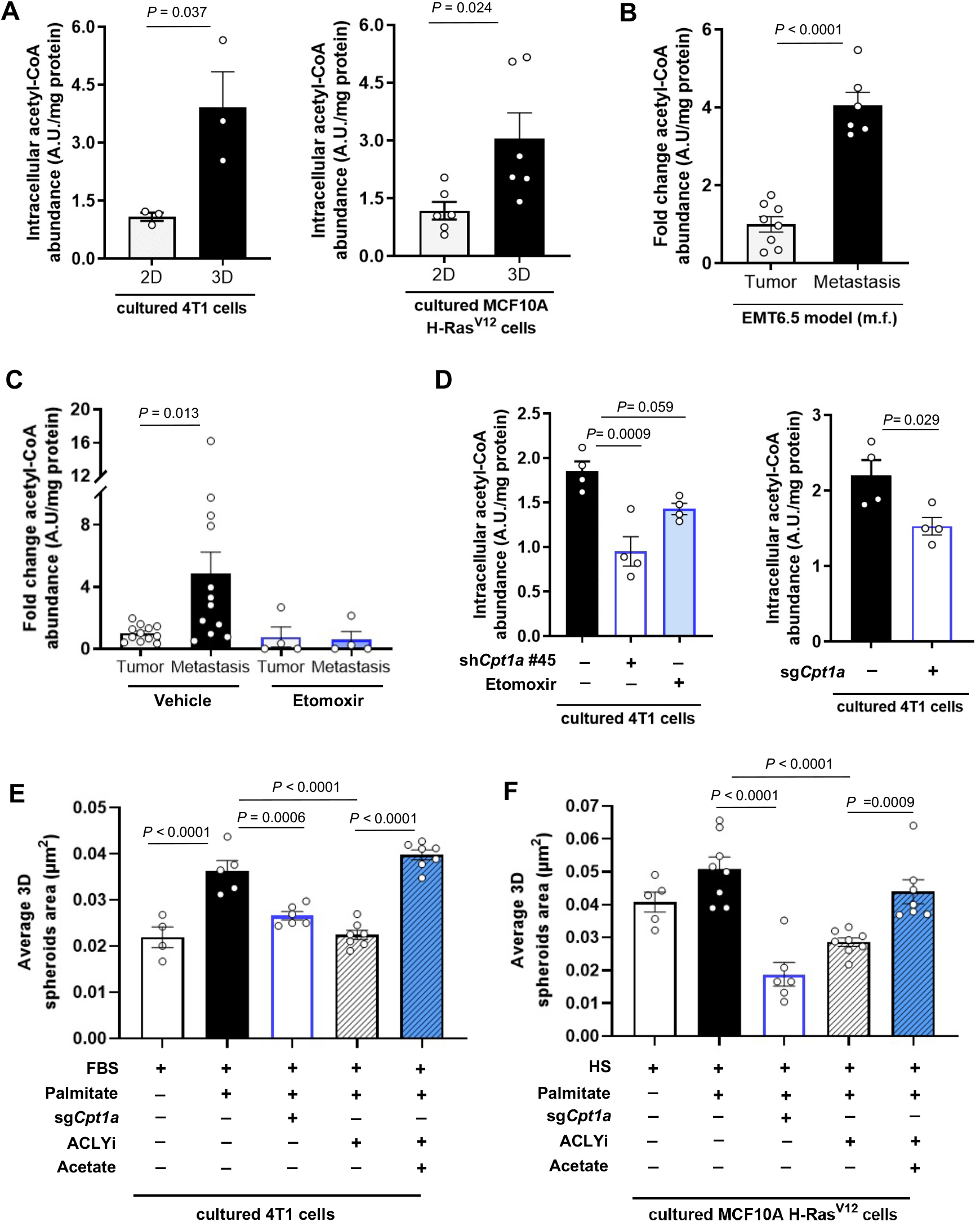
CPT1a activity sustains acetyl-CoA levels in spheroids and lung metastases. **A**. Intracellular levels of acetyl-CoA in 4T1 and MCF10A H-Ras^V12^ cells incubated in 2D monolayer and 3D spheroids cultures for 5 days in medium containing extra palmitate. Intracellular acetyl-CoA measurements in additional cell lines (EMT6.5 and MCF7) are shown in Supplemental Figure 4A. Two-tailed unpaired student’s T-test (n = 4). **B**. Relatives changes in acetyl-CoA abundance in EMT6.5 (m.f.) breast primary tumors and lung metastases. Data are shown as fold changes compared with the acetyl-CoA abundance in the primary tumors. Two-tailed unpaired student’s T-test (n = 4). **C**. Relatives changes in acetyl-CoA abundance in 4T1 (m.f.) breast primary tumors and lung metastases upon acute inhibition of CPT1A using the inhibitor etomoxir (30 mg/kg i.p.) or vehicle (water). Data are shown as fold changes compared with the acetyl-CoA abundance in the primary tumors of the group of mice treated with the vehicle. Data points represented as zero were below the detection limit. One-way ANOVA with Dunnett’s multiple comparison test (n ≥4). **D**. Relatives changes in acetyl-CoA abundance in mouse 4T1 breast cancer spheroids upon treatment with the CPT1A inhibitor etomoxir (50 μM), transduced with a lentiviral vector with shRNA against Cpt1a (knockdown), sgRNA Cpt1a (knockout) or scrambled control sequences in the presence of extra palmitate. Intracellular acetyl-CoA measurements in additional human cell lines (MCF10A H-Ras^V12^ and MCF7) are shown in Supplemental Figure 2B. One-way ANOVA with Dunnett’s multiple comparison test or two-tailed unpaired student’s T-test are shown (n = 4). **E** and **F**. 3D spheroids growth upon genetic inhibition of either *Cpt1a* or *CPT1A* together with pharmacologic ALCY inhibition using BMS-303141 (20 μM) in 4T1 (**A**.) and MCF10A H-Ras^V12^ (**B**) cells with or without extra palmitate and in the presence of the acetate as metabolic rescue (5 mM). 3D spheroid growth is represented by the average spheroids area of >100 spheroids. One-way ANOVA with Turkey’s multiple comparison test (n ≥4).

### CPT1a-mediated palmitate oxidation increases NF-*κ*B signaling

Next, we sought to identify how acetyl-CoA production by CPT1a supports metastasizing cancer cells. To address this question, we determined whether acetyl-CoA was required in the mitochondria or the cytosol. If acetyl-CoA is required in the cytosol, inhibiting ATP citrate lyase (ACLY) is expected to phenocopy CPT1a inhibition **(Supplemental Figure 4E)**. Indeed, we found that inhibiting ACLY with BMS-303141 (20μM, ACLYi) decreased 3D spheroid growth to a similar level to CPT1a inhibition and that acetate supplementation rescued this growth defect **(Figure 4E and F)**. This suggests that palmitate-derived acetyl-CoA was required in the cytosol, rather than the mitochondria, to induce spheroid growth. Accordingly, the level of ATP, a product of mitochondrial acetyl-CoA metabolism, was not significantly altered upon CPT1a deletion and acetate supplementation **(Supplemental Figure 4F)**. We then investigated possible fates of acetyl-CoA beyond mitochondria metabolism. One such possibility is histone acetylation ^45^. Yet, we found no prominent changes in histone acetylation in 3D cultured 4T1 cells upon CPT1a silencing using proteomics **(Supplemental Figure 4G)**. Based on these findings we hypothesized that palmitate-derived acetyl-CoA production may be important for non-histone protein acetylation, which may be connected to gene expression regulation ^46^. Hence, we investigated whether there were gene expression signatures that decreased upon CPT1a deletion, that were rescued by acetate, and that could explain the palmitate-promoted increase in spheroid growth. We performed RNA sequencing and consecutive gene set enrichment analysis (GSEA) in 4T1 cells in these conditions. Among the highest-ranking hallmark gene sets, we observed a signature for NF-κB signaling that was decreased upon CPT1a deletion and rescued upon acetate treatment **(Figure 5A and Supplemental Figure 5A)**. Key genes of this signature were confirmed in an additional breast cancer cell line **(Supplemental Figure 5B)**. Conversely, we analyzed all CPT1a-dependent and acetate rescued gene expression changes to predict upstream regulators using Ingenuity Pathway Analysis ^47^. Remarkably, several upstream regulators of the NF-κB signaling pathway (*Nfkbia, Chuk, Ikbkb, Tnf, Traf3ip2, Igf1, Tnsf11, Rela/p65* and *Ikbkg*) scored in the top 30 of the predicted activated regulators **(Figure 5B)**. Thus, this shows that NF-κB signaling is likely induced by palmitate in a CPT1a-dependent manner. NF-κB signaling is known to induce pro-metastatic gene expression programs such as an epithelial-to-mesenchymal transition (EMT) ^1,48^. Accordingly, we found a reduced EMT signature upon CPT1a deletion that was rescued with acetate supplementation **(Supplemental Figure 5C)**. Thus, we concluded that NF-κB signaling is activated downstream of the CPT1a-mediated palmitate oxidation in 3D spheroids.

**Figure 5.**
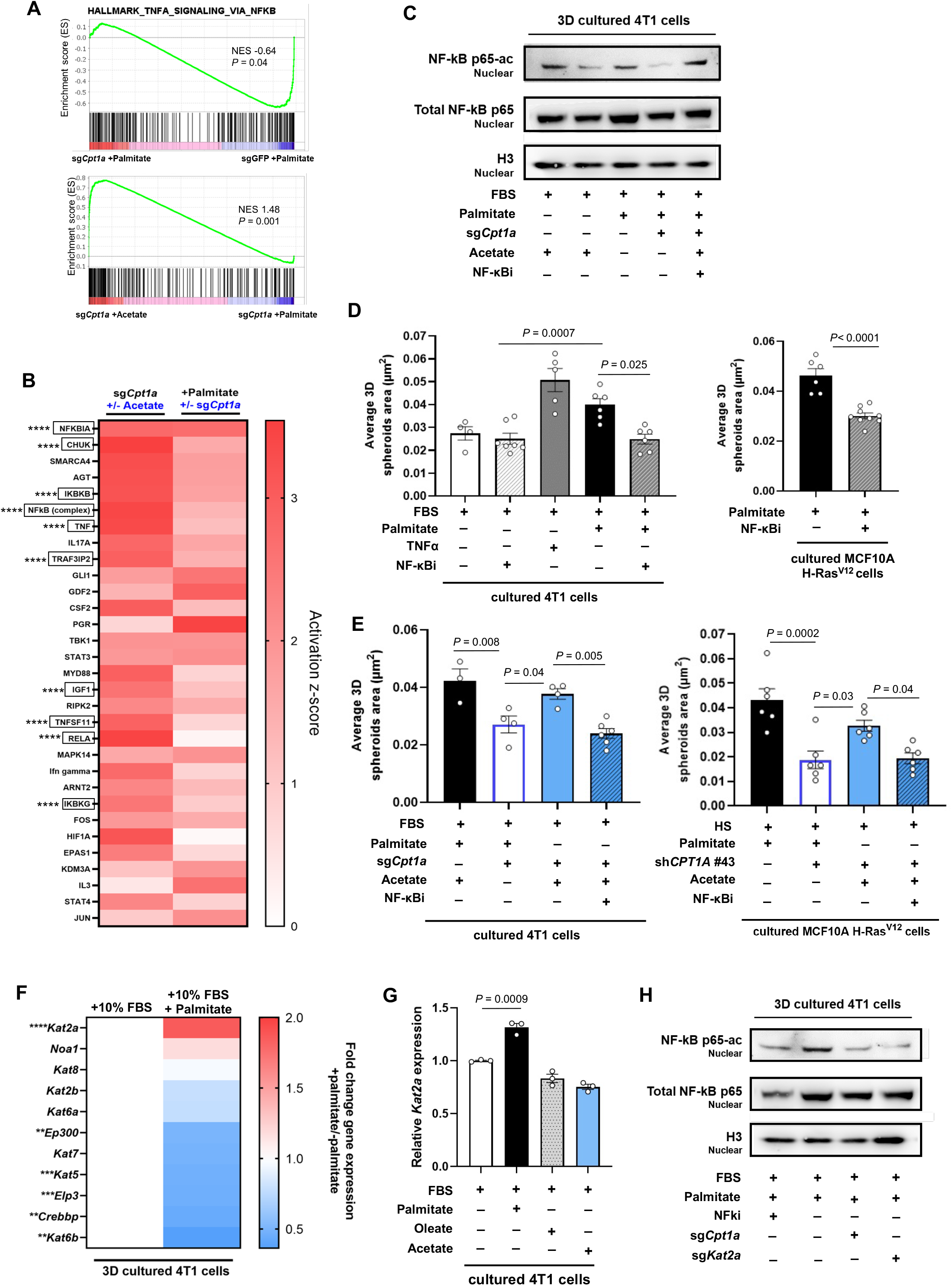
CPT1a activity is required for p65 acetylation and NF-κB signaling. **A**. GSEA enrichment plots comparing the gene expression profiles in 4T1 3D spheroids transduced with a lentiviral vector containing sg*Cpt1a* or control sgGFP (top panel) and sg*Cpt1*a 4T1 3D spheroids cultured with or without acetate (bottom panel). NES, normalized enrichment score; the P value indicates the significance of the enrichment score. **B**. Upstream regulator analysis performed using Ingenuity Pathway Analysis using the differential gene expression of CPT1A knockout (*sgCpt1a*) 4T1 3D spheroids in the presence or absence of acetate (metabolic rescue, 5 mM) as input. Activation score of the top 30 upstream regulators (left column) was compared to those predicted for the differential gene expression of CPT1A knockout versus control conditions (in the presence of extra palmitate). Genes related to the activation of the NF-κB pathway are framed. Asterisks represent the overlap p-value calculated using Fisher’s Exact Test (****p < 0.0001) **C**. Acetylated p65 (NF-κB p65ac) in nuclear extracts of 3D spheroids transduced with a lentiviral vector containing sg*Cpt1a* or control sgGFP and cultured for 3 days in the presence of extra palmitate (75 μM), acetate (5 mM) or the NF-κB inhibitor PDTC (0.5 μM). Histone H3 is shown as a loading control. A representative of three experiments is shown. **D**. 3D growth of 4T1 (left panel) and MCF10A H-Ras^V12^ (right panel) cells upon treatment with the NF-κB inhibitor (PDTC, 0.5 μM) in the presence of extra palmitate and 4T1 3D spheroids growth upon activation of the pathway via either supplementation of TNFa (10 ng/μL) or extra palmitate (75 μM) is shown. 3D spheroid growth is represented by the average spheroids area of >100 spheroids. One-way ANOVA with Turkey’s multiple comparison test (4T1, n ≥4) and two-tailed unpaired student’s T-test (MCF10A H-Ras^V12^, n = 4). **E**. 3D growth of 4T1 (left panel) and MCF10A H-Ras^V12^ (right pale) cells upon treatment with the NF-κB inhibitor to the inhibitory impact of CPT1a inhibition (sg*Cpt1a*) in the presence of the extra palmitate (75 μM) or acetate as metabolic rescue (5 mM). 3D spheroid growth is represented by the average spheroids area of >100 spheroids. One-way ANOVA with Turkey’s multiple comparison test (n ≥4). **F**. Differential expression of acetyltransferases in 4T1 3D spheroids in the presence or absence of extra palmitate in culture medium after 5 days of incubation. Data represent the fold changes compared with the non-extra palmitate condition. Two-tailed unpaired student’s T-test (n = 4). Asterisks represent statistical significance as follows: *p < 0.01, ***p < 0.001, ****p < 0.0001. **G**. Relative *Kat2a* expression in 4T1 3D spheroids with or without additional palmitate (75 μM), oleate (116 μM) or acetate (5 mM). One-way ANOVA with Turkey’s multiple comparison test (n =6). **H**. Acetylated p65 (NF-κB p65ac) in nuclear extracts of 3D spheroids transduced with a lentiviral vector containing sg*Cpt1a*, sg*Kat2a* or control sgGFP, and cultured for 3 days in the presence of extra palmitate (75 μM) or the NF-κB inhibitor PDTC (0.5 μM). Histone H3 is shown as a loading control. A representative of three experiments is shown.

### CPT1a activity induces NF-κB signaling by promoting acetylation of the NF-*κ*B subunit p65

Interestingly, acetylation of NF-κB family members is known to dynamically regulate NF-κB activation and transcriptional activity ^49^. Thus, we asked whether p65 acetylation is regulated by CPT1a. To address this question, we determined the amount of total p65 binding to the DNA by electrophoretic mobility shift assay (EMSA) and p65 acetylated at K310 in the nucleus upon CPT1a silencing and upon acetate supplementation in 4T1 cells using acetylation-specific western blot analysis. Total nuclear p65 and DNA binding activity were not altered **(Figure 5C, Supplemental Figure 5D)**, which may indicate that the genome-wide distribution of p65 is not altered. Strikingly, however we found that K310 acetylated p65 in the nucleus decreased upon CPT1a silencing and was rescued upon acetate supplementation **(Figure 5C)**. These findings suggest an important and specific role of p65 acetylation in mediating the induction of distinct NF-κB targets.

Next, we decided to modulate NF-κB signaling and assessed 3D spheroid growth. TNFα is an activator of NF-κB, while pyrrolidinedithiocarbamate ammonium (PDTC, 0.5 μM abbreviated also as NF-κBi) blocks the translocation of p65 into the nucleus and hence reduces its transcriptional activity ^50^. Consistent with the importance of palmitate-induced acetylation of p65 for spheroid growth, treating 4T1 and MCF10A H-Ras^V12^ cancer cells with TNFα increased spheroid growth even in the absence of palmitate, while PDTC (NF-κBi) treatment decreased spheroid growth in the presence of palmitate to the same level as without extra palmitate **(Figure 5D)**. Accordingly, acetate supplementation did not any longer rescue spheroid growth in the presence of PDTC (NF-κBi) **(Figure 5E)**. Therefore, we concluded that palmitate-derived acetyl-CoA production is required for p65 acetylation and downstream NF-κB signaling.

### KAT2a acetylates p65 in the presence of palmitate and is required for palmitate-promoted spheroid growth

Acetyl-CoA is a global product of fatty acid oxidation. Thus, we asked how palmitate, but not other fatty acids, can increase p65 acetylation. One possibility is that a specific acetyltransferase mediates the palmitate-derived acetylation of p65. To address this hypothesis, we compared the gene expression of several acetyltransferases in 3D cultured 4T1 cells in the presence and absence of extra palmitate. We found that only one acetyltransferase, namely lysine acetyltransferase 2a (KAT2a or GCN5) was highly induced in the presence of palmitate **(Figure 5F)**. Strikingly, neither oleate nor acetate addition did increase KAT2a expression **(Figure 5G)** although both increase acetyl-CoA levels **(Supplemental Figure 6A)**. This observation suggests that specificity may be achieved through the expression of KAT2a.

KAT2a has been mainly studied concerning histone H3 acetylation and in particular H3K9 acetylation as well as some non-histone targets^51-53^. Therefore, we evaluated whether KAT2a could affect p65 indirectly via H3K9 acetylation by ChIP-sequencing. We found that the loss of either KAT2a or CPT1a only resulted in minor changes in H3K9-acetylation **(Supplemental Figure 6B)**. These minor changes in H3K9-acetylation after CPT1a deletion were not rescued by acetate **(Supplemental Figure 6C)**. Despite that, a clear decrease in the transcription of NF-κB targets upon CPT1a or KAT2a deletion and rescue with acetate was observed **(Supplemental Figure 6D)**. Consequently, we deemed it unlikely that KAT2a acts to promote metastasis via histone acetylation.

On the other hand, it has been shown that KAT2a physically interacts with p65 in vivo, which activates gene expression via acetylation of the p65 subunit during hippocampal memory regulation ^54^. Thus, we hypothesized that KAT2a modulates gene expression of pro-metastatic NF-κB targets by mediating directly p65-K310 acetylation. To address this hypothesis, we deleted KAT2a in 4T1 cells and assessed the presence of K310 acetylated p65 in the nucleus. As observed with CPT1a deletion, total p65 and general binding to DNA were not significantly altered **(Figure 5H, Supplemental Figure 5D)**. Yet, in line with the hypothesis that KAT2a is responsible for acetylating p65 in the presence of palmitate, we observed that its deletion decreased specifically the amount of p65-K310ac in the nucleus **(Figure 5H)**. Subsequently, we investigated whether blocking KAT2a and thus p65 acetylation, prevents the palmitate-induced spheroid growth. Like CPT1a also KAT2a inhibition did not decrease proliferation of cancer cells growing in 2D monolayer **(Supplemental Figure 6E)**. However, we found that KAT2a deletion reduced 4T1 spheroid growth to a similar degree as CPT1a **(Figure 6A, Supplemental Figure 6F**). Accordingly, 3D spheroid growth of 4T1, MCF7 and MCF10A H-Ras^V12^ was significantly reduced upon treatment with cyclopentylidene-[4-(4’-chlorophenyl)thiazol-2-yl)hydrazone (CPTH2), an inhibitor of KAT2a (2 μM) ^55^ (Figure 6A and B). Moreover, breast cancer cells showed more sensitivity to CPTH2 in the presence of extra palmitate compared to the absence of extra palmitate **(Supplemental Figure 6G)**. Based on these data we concluded that KAT2a acetylates p65 at K310 in the presence of palmitate and that this activity is essential for palmitate-promoted 3D growth.

**Figure 6.**
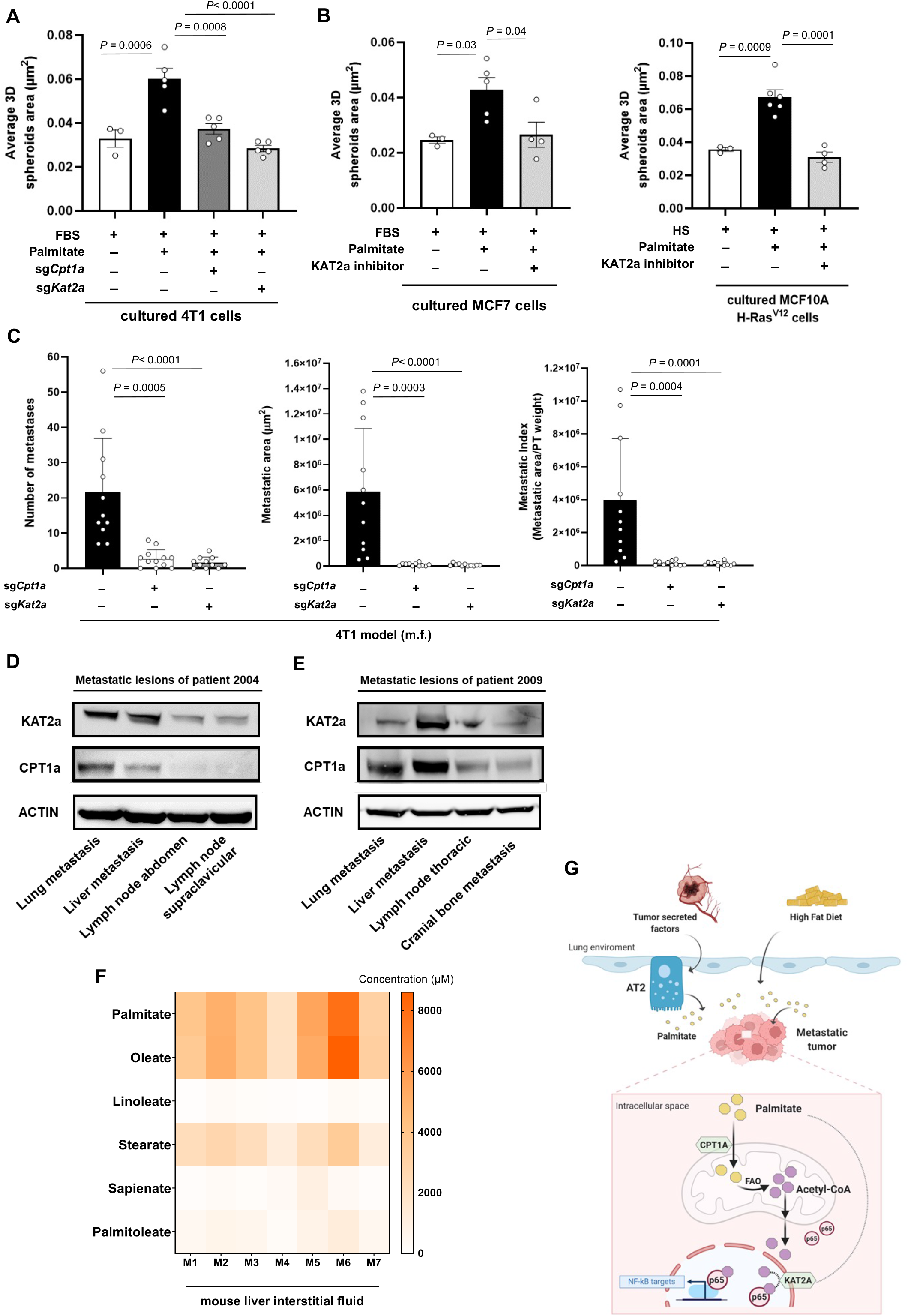
Silencing CPT1a and KAT2a inhibit metastasis formation in vivo and are co-expressed in palmitate-enriched environments. **A**. 3D spheroid growth of 4T1 cells silenced for *Cpt1a* or *Kat2a* with or without extra palmitate. One-way ANOVA with Turkey’s multiple comparison test (n ≥4). **B**. 3D spheroid growth in MCF7 and MCF10A H-Ras^V12^ cells upon pharmacologic inhibition of KAT2a using the inhibitor CPTH2 (5μM) cultured for 5 days in medium with or without extra palmitate supplementation. 3D spheroid growth is represented by the average spheroids area of >100 spheroids. One-way ANOVA with Turkey’s multiple comparison test (n ≥4). **C**. Metastatic burden in mice injected with 4T1 (m.f.) lungs upon genetic knockout of *Cpt1a* and *Kat2a*. m.f.: mammary fat pad; i.v.: intravenous. Data for control and sg*Cpt1a* are also shown in Figure 3E. One-way ANOVA with Dunnett ‘s multiple comparison test (n ≥11). **D** and **E**. CPT1a and KAT2a protein expression in different metastasis sites of breast cancer patient 2004 (D) and 2009 (E) from the UPTIDER program. **F**. Fatty acid concentration present in the liver interstitial fluid, detected by mass spectrometry from healthy BALB/c mice (n =7). **G**. Tumor secreted factors and high fat diet are two independent factors increasing palmitate levels in the lung environment. Metastasizing cancer cells use the available palmitate to drive p65 acetylation in a CPT1a dependent manner resulting in pro-metastatic NF-κB signaling.

### KAT2a inhibition impairs metastasis formation in mice and KAT2a is co-expressed with CPT1a in metastases from patients with breast cancer

Next, we assessed whether blocking KAT2a and hence p65 acetylation is sufficient to impair lung metastasis formation to a similar extent than CPT1a deletion. We injected 4T1 control and KAT2a knockout cells to the mammary fat pad of mice and assessed lung metastases number, area and metastatic index (metastases area divided by primary tumor weight) based on H&E staining. Strikingly, we observed that metastases number, metastases area and metastatic index were highly reduced in the absence of KAT2a expression compared to control **(Figure 6C, Supplemental Figure 3A)**, while primary tumor growth showed only a small reduction compared to control **(Supplemental Figure 3B)**. Thus, we concluded that blocking KAT2a phenocopies CPT1a inhibition and is sufficient to impair metastatic growth in mice.

Finally, we investigated whether there is evidence of this CPT1a-KAT2a-driven mechanism in metastatic patients with breast cancer. In this case we expect that metastases growing in palmitate-enriched organs require co-expression of CPT1a and KAT2a. We therefore collected several metastatic lesions of different organs from two patients with breast cancer that had agreed to be part of the UPTIDER rapid autopsy program at UZ Leuven (see **Supplemental Table 3** for clinicopathological information) and determined CPT1a and KAT2a protein expression based on western blot analysis. Strikingly, we observed that CPT1a and KAT2a proteins were co-expressed with the highest expression in lung and liver metastases compared to cranial bone and lymph node metastases **(Figure 6D and E)**. These data are consistent with our *in vitro* and in mice determined mechanism requiring simultaneous activity of CPT1a and KAT2a for metastases growth in palmitate-enriched environments such as the lung and liver **(Figure 1A and 6F)** compared to other environments such as lymph node ―which has been found to be oleate enriched instead ^18^. We therefore concluded that CPT1a and KAT2a may act in concert to promote metastatic growth specifically in palmitate-enriched organ environments in patients with breast cancer.

In conclusion, we identify the lung as palmitate-rich environment whose response to pathological conditions of pre-metastatic niche formation and obesity converges into an even further increase in palmitate availability. Metastasizing cancer cells rely on CPT1a-dependent palmitate oxidation to acetyl-CoA, which in turn serves as a substrate for the acetylation of p65 by KAT2a. Subsequently the pro-metastatic NF-κB transcriptional program is activated supporting metastatic growth in palmitate-rich environments **(Figure 6G)**.

## Discussion

Here, we provide the first analysis of lipid availability in healthy and pathological conditions of the lung which is a frequent metastatic site across many cancers. Furthermore, we determine a novel mechanism linking palmitate-rich environments to protein acetylation upstream of pro-metastatic NF-κB signaling.

Palmitate has been recently shown to induce a pro-metastatic memory in primary tumors associated with the stable deposition of histone H3 lysine 4 trimethylation by the methyltransferase Set1A and a consequent neural signature that stimulates intratumor Schwann cells and innervation ^17^. Here, we identify palmitate as an important nutrient for cancer cells growing in distant organs, and we describe an epigenetic independent role of palmitate in promoting metastasis formation.

In macrophages, endothelial cells and adipocytes ^56-59^ palmitate, like obesity ^60^, has been associated with inflammatory NF-κB signaling. However, the underlying mechanism and whether this occurs in cancer and metastasis formation is largely unknown with only one report linking etomoxir treatment to reduced phosphorylation of IκBα ^61^. Here, we discover a mechanism by which CPT1a-dependent palmitate oxidation regulates NF-κB signaling via acetylation of p65. Moreover, we find that the acetyltransferase KAT2a, which was previously not linked to p65 acetylation in cancer^62^, is induced by palmitate to acetylate p65. This provides for the first-time evidence of a nutrient environment-dependent regulation of protein acetylation in organs of metastasis.

Acetyl-CoA levels have been previously linked to protein acetylation of Smad induced by leptin signaling ^63^ and histone acetylation ^64-66^ with subsequent breast cancer metastasis ^63,64^. We add to this current knowledge by defining p65 as an acetylation target modulated by palmitate-derived acetyl-CoA. Contrary to a previous study showing that KAT2a participates independent of its acetyltransferase activity in p65/RelA degradation ^67^, and in line with the previously reported interaction of KAT2a and p65 activating gene expression via acetylation ^54^ we find that KAT2a activity is needed to activate NF-κB signaling in palmitate rich environments.

Several roles of fatty acid oxidation have been described in metastasizing cancer cells ^1,12^. In breast cancer cells, CPT1a deletion has been shown to inhibit lymph angiogenesis ^68^ and a splice variant of CPT1a has been found to promote histone deacetylase (HDAC) activity in the nucleus through a protein–protein interaction ^69^. Moreover, ATP levels sustained by CPT1a activity have been shown to enable the activity of SRC oncoprotein through autophosphorylation ^70^. Beyond breast cancer, in matrix detached colorectal cancer cells ROS accumulated upon CPT1a silencing ^71^ and in matrix detached ovarian cancer cells oleate was oxidized to sustain mitochondrial respiration ^72^. Similarly, oleate enriched lymph node metastases from melanoma relied on fatty acid oxidation ^73^. We add to these various functions of fatty acid oxidation by showing that palmitate oxidation-derived acetyl-CoA is important for p65 acetylation. Moreover, we show that CPT1a silencing is sufficient to target metastasis formation in lean and obese mice, the latter has not been shown *in vivo* before. This may suggest that the use of the CPT1a inhibitor etomoxir should be revisited especially for obese breast cancer patients.

To date, very little is known about the priming of the pre-metastatic niche by primary tumor-secreted factors with only one report highlighting spared glucose consumption through the induction of a hypometabolism of resident cells ^21^. We discover that, mimicking the systemic effect of a primary tumor by inducing pre-metastatic niche formation, only palmitate availability is selectively increased. Furthermore, we add to the sparse knowledge on nutrient priming of the pre-metastatic niche by showing that palmitate is actively secreted by lung resident AT2 cells in response to tumor secreted factors. Moreover, we find that high-fat diet alters the lung environment by increasing the availability of selected fatty acids in the interstitial fluid, which may be reflective of the systemic increase that has been observed in circulation ^41^. Obesity is a condition highly prevalent in breast cancer patients and multiple mechanisms linking signaling for example through leptin and insulin to cancer progression have been defined. However, we show for the first time that obesity resulting from high fat diet feeding increases intra-lung fatty acid availability even though lungs are non-steatotic organs. Thus, this suggests a much broader impact of diet on nutrient availability than previously considered and highlights an underappreciated role of nutrient pre-conditioning of distant organs during metastasis formation.

## Acknowledgments

We thank Peter Carmeliet (VIB-KU Leuven) for providing short hairpin RNA (shRNA) against *Cpt1a*, David Nittner (VIB Histology and Imaging Core) for their histology services. We would like to thank Theo Killian and Vincent van Hoef (VIB Bioinformatics Core Facility) for their help with the RNAseq analysis, and Prof. László Tora for his advice in ChIPseq analysis. We want to acknowledge all physicians from the multidisciplinary breast centre Leuven (especially Patrick Neven, Ann Smeets, Ines Nevelsteen, Kevin Punie) for their clinical contribution to the UPTIDER and/or CHEMOREL cohorts. The authors also want to acknowledge the patients who accepted to donate tissues post-mortem in the context of the UPTIDER program, as well as the whole multidisciplinary UPTIDER team.

## Author’s contributions

Conception and design: PAM, GD and S-MF.

Development of methodology: PAM, GD, AC, MP, and CRD.

Acquisition of data (including providing animals, acquiring and managing patient samples, providing facilities, etc.): PAM, GD, AC, EN, AM, MP, JVE, IV, DB, TG, MDS, SL, FR, SH, YL, QW, EK, CCA, VG, GF, J-CM, DL, MM, VP, SRZ, JC, HW, VB, CD, TGPG and IM.

Analysis and interpretation of data (e.g., statistical analysis, biostatistics, computational analysis): PAM, JFG, FCA, MP, OM-B, QW, FR, BB, SD, AA, VP, and S-MF.

Writing, review, and/or revision of the manuscript: PAM and S-MF.

Administrative, technical, or material support (i.e., reporting or organizing data, constructing databases): PAM, DB, MP and JVE.

Study supervision: S-MF.

---

**Supplemental Figure 1.**
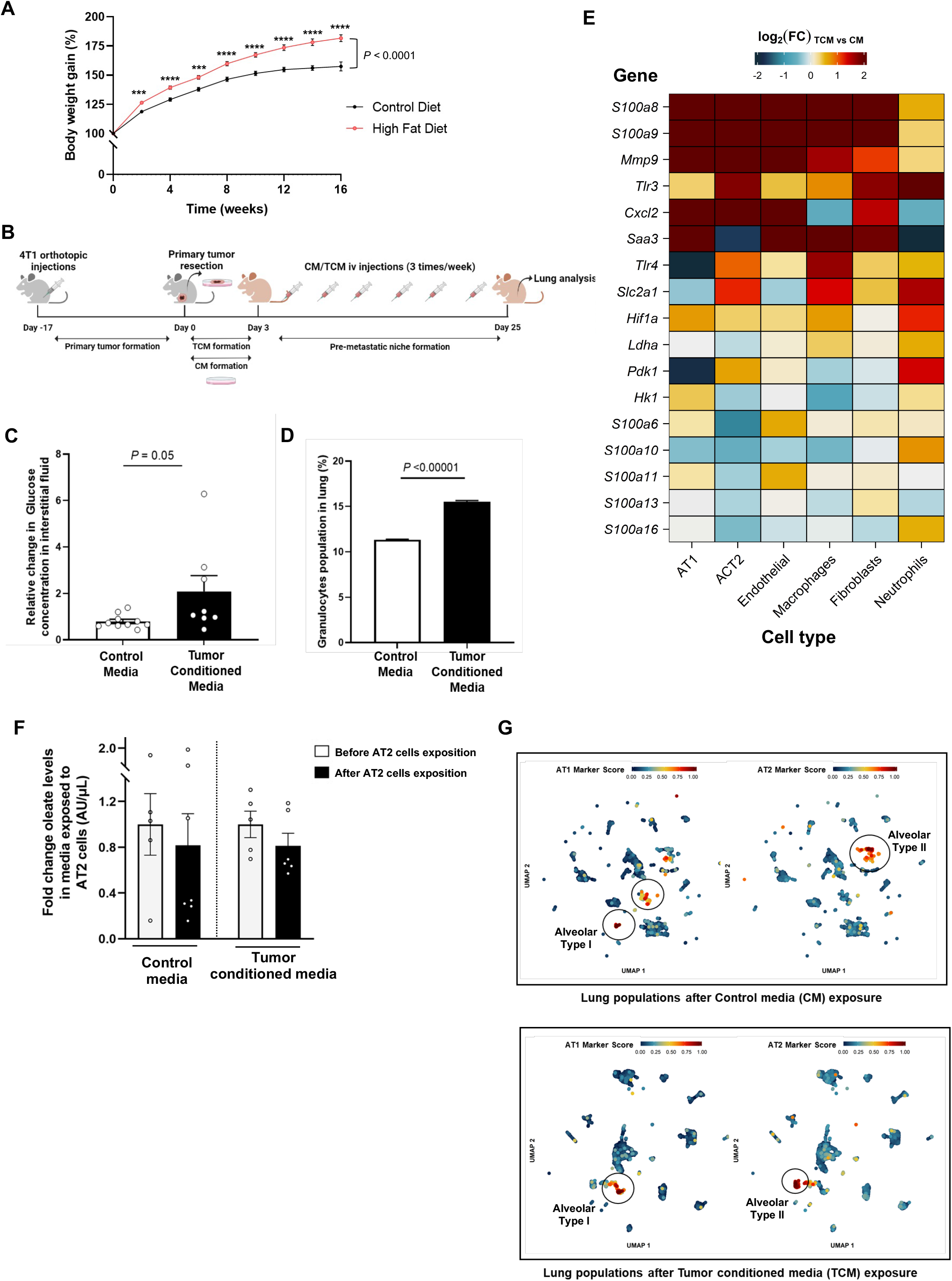
Palmitate is released by AT2 cells during pre-metastatic niche formation. **A**. Mouse weight gain upon high fat or control diet over the course of the experiment (n =60). Mixed-effects analysis with Sidak’s multiple comparisons. Asterisks represent statistical significance as follows: ***p < 0.001, ****p < 0.0001. **B**. Schematic illustration for experimental pre-metastatic niche formation procedure. CM, control media; TCM, tumor conditioned media; i.v intravenous. **C**. Relative glucose concentration in the lung interstitial fluid of healthy BALB/c mice exposed to control media or tumor condition media. Two-tailed unpaired student’s T-test (n ≥7). **D**. Granulocytes population (which includes neutrophils) present in lungs after induction of pre-metastatic niche formation using tumor conditioned media or control media. Two-tailed unpaired student’s T-test (n =4). **E**. Changes in gene expression in lung populations after tumor conditioned media or control media injections, that have been previously linked to pre-metastatic niche formation, such as increased expression of *S100a8, S100a9* and *Mmp9* genes ^31,34^ *Tlr3* activation and *Cxcl2* production in lung alveolar type II cells ^31^, *Tlr4* and *Saa3* overexpression in lung endothelial cells and macrophages ^32^, increased expression of *Slc2a1, Hif1a*, and *Ldha* in macrophages ^33^ as well as upregulated S100 genes in lung fibroblasts ^35^. Scaled expression levels are indicated by the color scale, where log2 FC denotes fold change TCM vs. CM. **F**. Relative oleate levels present in control or tumor conditioned media before and after exposure to lung resident AT2 cells for 72h. Data are shown as fold changes compared with oleate abundance in media before incubation with AT2 cells. Two-tailed unpaired student’s T-test (n =6). **G**. UMAP plots for the scRNA-seq data corresponding to lungs preconditioned with control media (CM) and tumor conditioned media (TCM). Color-coded based on GSVA-based marker scores for gene sets corresponding to alveolar type I (AT1) and II (AT2) marker genes. Identified clusters are indicated within black circles. Marker scores are scaled to the range 0–1 for each individual market set (see Methods).

**Supplemental Figure 2.**
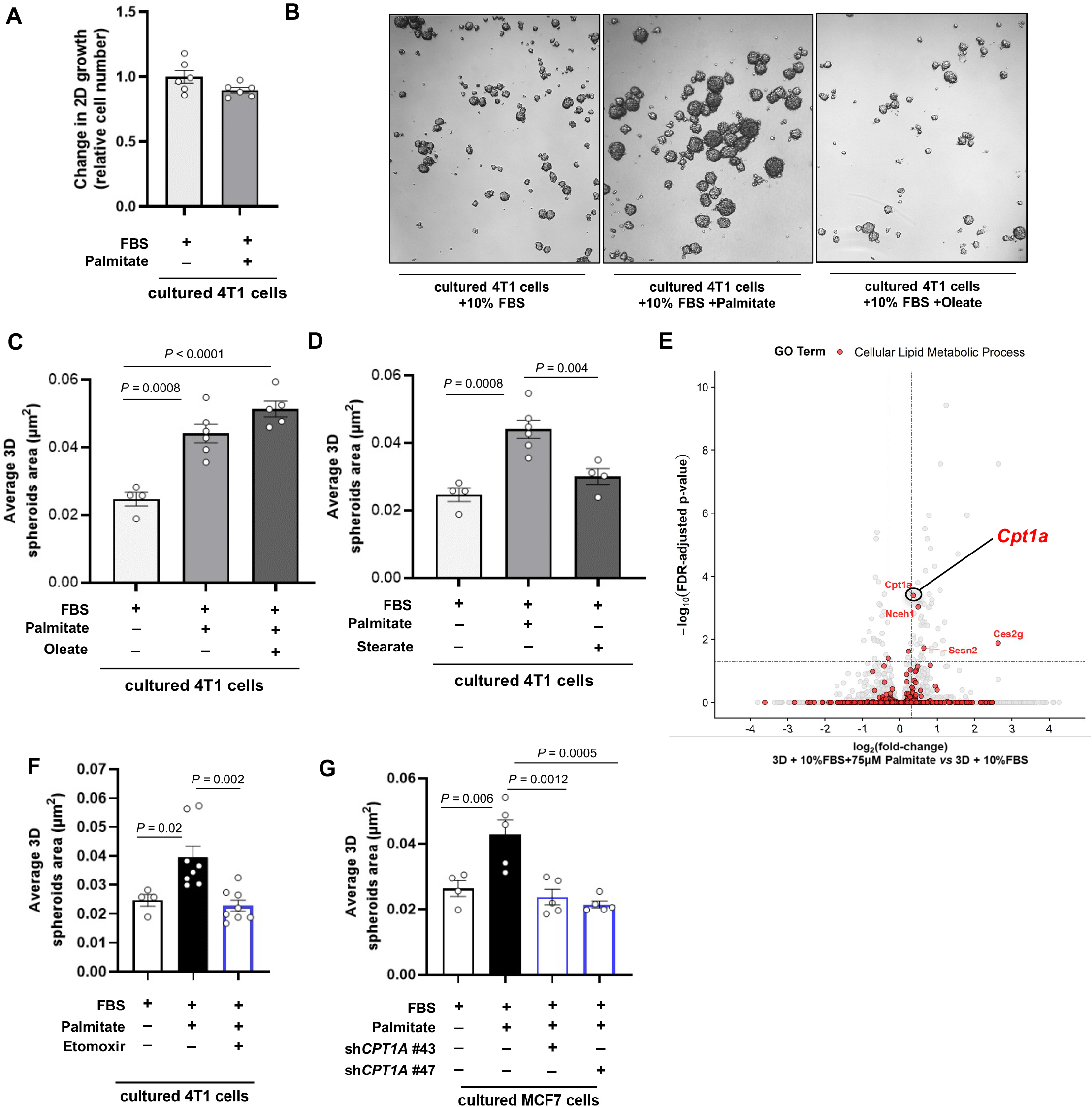
Oleate and stearate do not phenocopy palmitate response. **A**. Relative proliferation of 4T1 cells (with or without extra palmitate) normalized to condition without extra palmitate. Two-tailed unpaired student’s T-test (n =6). **B**. Representative pictures of 4T1 spheroids cultured in 10%FBS, or 10%FBS in the presence of palmitate (75 μM) or oleate (116 μM). **C** and **D**. 3D spheroids growth of 4T1 cells upon palmitate plus oleate supplementation (**C**) and stearate (**D**) supplementation represented by the average spheroids area of >100 spheroids. One-way ANOVA with Holm-Sidak’s multiple comparison test (n=≥4). **E**. Differentially expressed genes in 3D spheroid 4T1 in the presence or absence of extra palmitate (10% FBS + palmitate or 10% FBS). Calculated differences of gene expression are presented by plotting the negative log10 of false discovery rate (Y-axis) against the log2 fold change of gene expression (X-axis). In red, genes belong to the GEO term lipid metabolic process (GO:000662). Only genes above the stablished cutoff are annotated. **F**. 3D spheroids growth of 4T1 cells upon CPT1a inhibition using etomoxir (50 μM) in the presence of extra palmitate (75 μM) represented by the average of spheroids area of >100 spheroids. One-way ANOVA with Turkey’s multiple comparison test (n = 4). **G**. 3D spheroids growth of MCF7 cells upon CPT1a knockdown compared to scrambled control upon palmitate supplementation (75 μM) represented by the average of spheroids area of >100 spheroids. One-way ANOVA with Turkey’s multiple comparison test (n = 4).

**Supplemental Figure 3.**
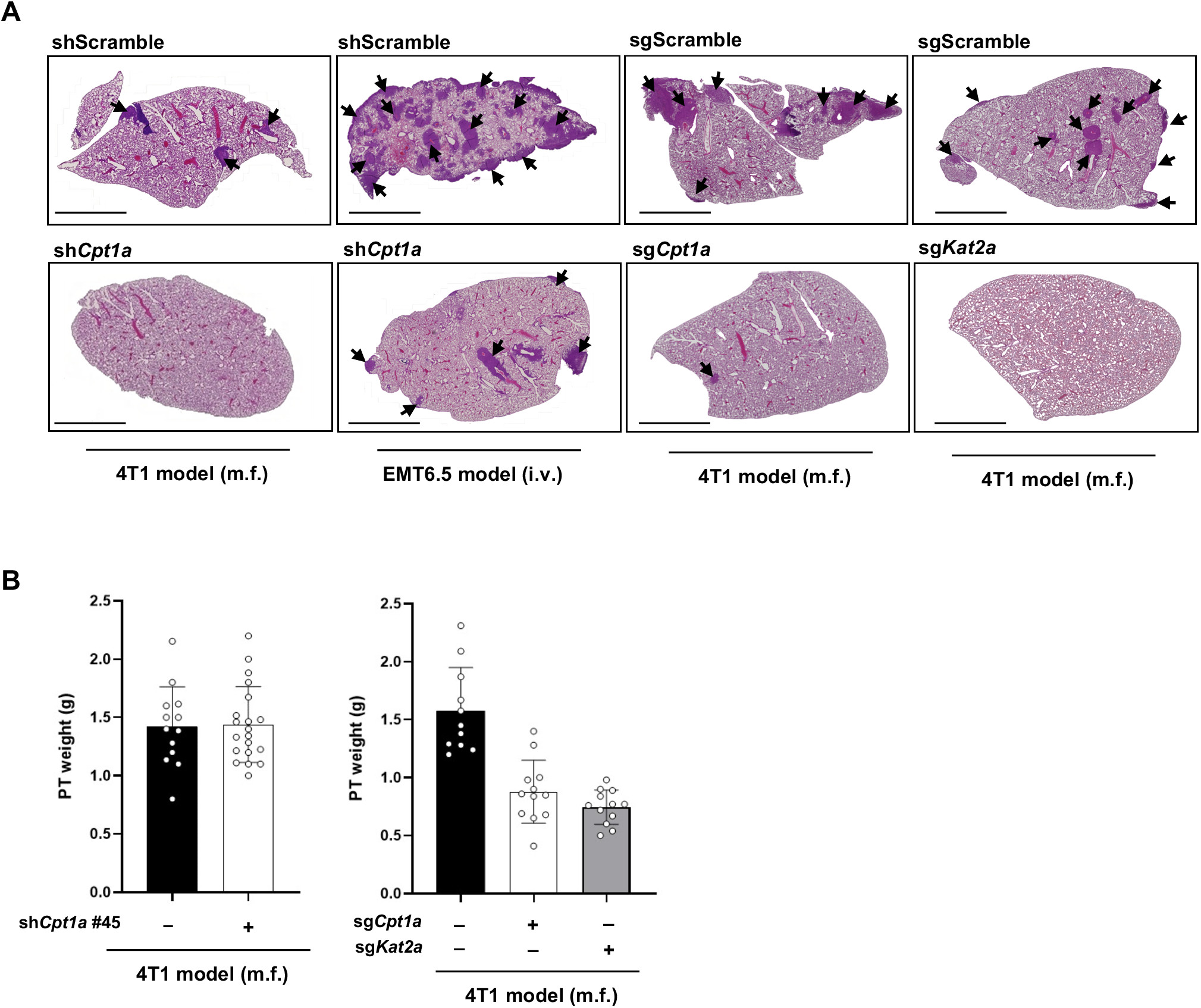
Primary tumor weight and lung H&E staining upon CPT1a and KAT2a loss. **A**. Representative pictures of tissue from the lung of mice injected with 4T1 (m.f.) and EMT6.5 (i.v) upon genetic inhibition of CPT1a based on H&E staining. Arrowheads indicate metastasis tissue. Scale bars, 0.5 cm. **B**. Final primary tumor weight upon genetic inhibition of CPT1A and KAT2A in the 4T1 model (m.f.). One-way ANOVA with Dunnett’s multiple comparison test (n ≥11).

**Supplemental Figure 4.**
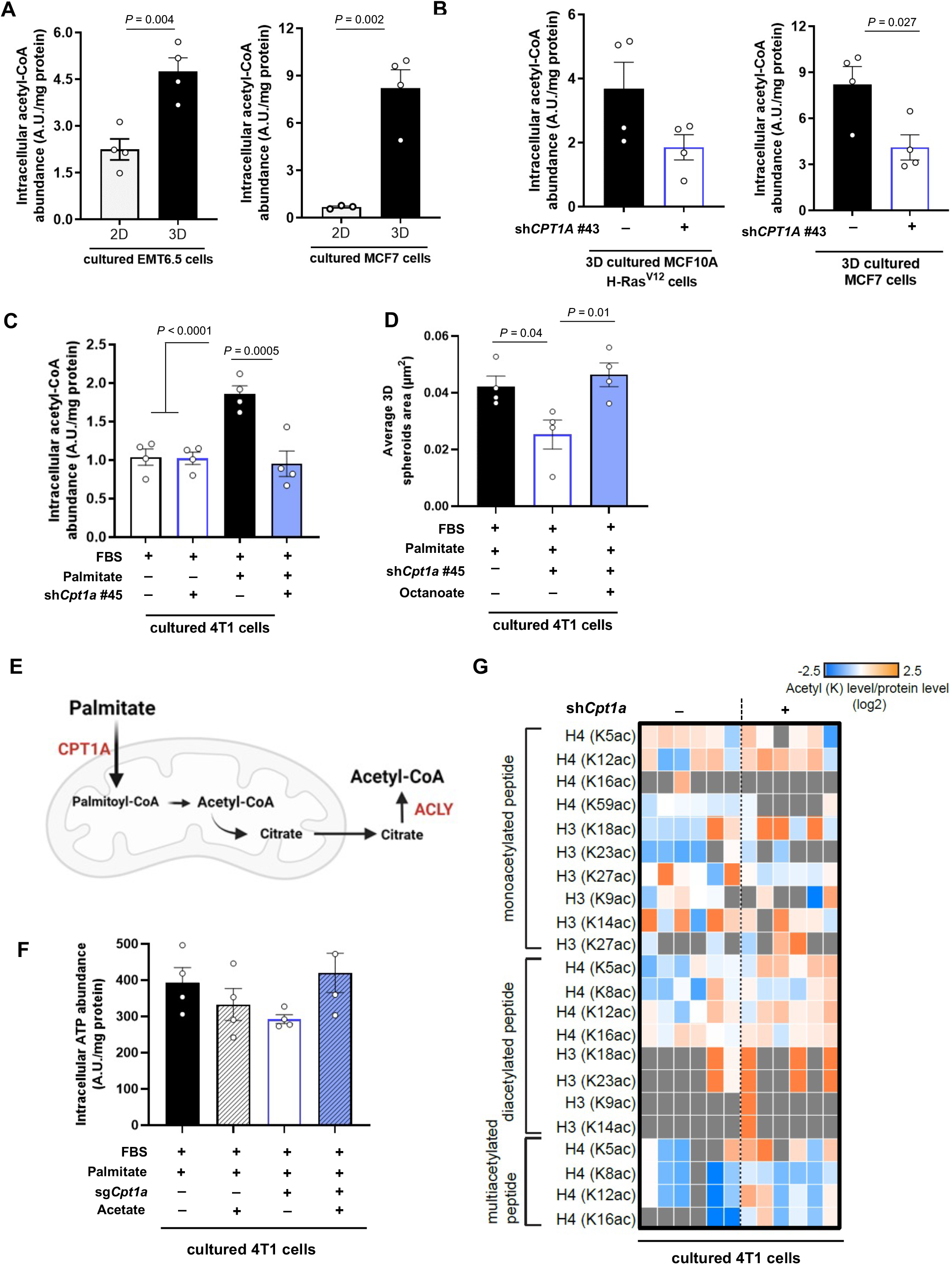
Breast cancer spheroids in the presence of additional palmitate rely on CPT1a for acetyl-CoA production and for sustaining palmitate-induced 3D growth. **A**. Intracellular levels of acetyl-CoA in EMT6.5 and MCF7 cells incubated in 2D monolayer and 3D spheroids cultures for 5 days in medium containing extra palmitate. Two-tailed unpaired student’s T-test (n = 4). **B**. Relatives changes in acetyl-CoA abundance in human MCF10A H-Ras^V12^ and MCF7 breast cancer spheroids transduced with a lentiviral vector with shRNA against *CPT1A* (knockdown) or scrambled control sequences in the presence of extra palmitate. Two-tailed unpaired student’s T-test is shown (n = 4). **C**. Relatives changes in acetyl-CoA abundance in mouse 4T1 breast cancer spheroids transduced with a lentiviral vector with shRNA against *Cpt1a* (knockdown) or scrambled control sequences in the presence or absence of extra palmitate. One-way ANOVA with Dunnett’s multiple comparison test (n = 4). **D**. 3D spheroids growth of 4T1 cells upon palmitate supplementation (75 μM), CPT1a genetic inhibition (sh*Cpt1a*) and metabolic rescue with octanoate (130 μM) represented by the average of spheroids area of >100 spheroids. One-way ANOVA with Dunnett’s multiple comparison test (n = 4). **E**. Schematic representation of the palmitate flux into the mitochondria via CPT1A and the ACLY-dependent export of the mitochondrial acetyl-Coa pool to the cytosol via citrate. **F**. Intracellular levels of ATP in CPT1a knockout and control 4T1 3D spheroids cultured for 5 days in medium containing extra palmitate (75 μM) or acetate as metabolic rescue (5 mM). **G**. Heatmap display of the log2 transformed ratios obtained for the indicated histone acetylation for CPT1a knockdown and control 4T1 3D spheroids cultured for 5 days in medium containing extra palmitate (75 μM). Relative abundances ratios, light/SILAC heavy, were obtained with the SILAC (Stable Isotope Labeling with Amino acids in Cell culture) internal standard strategy ^74^.

**Supplemental Figure 5.**
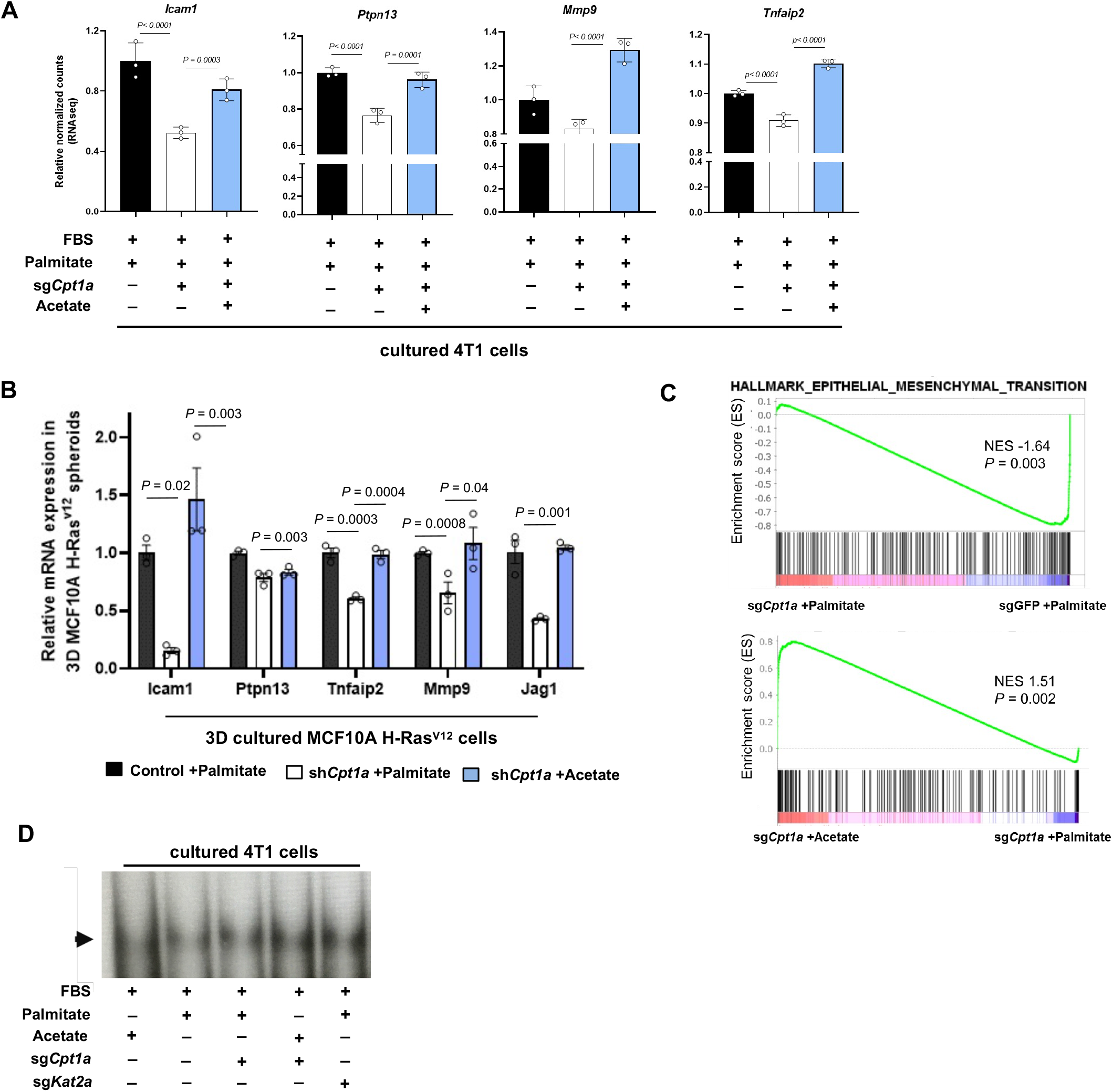
CPT1a deletion reduces NF-κB signaling pathway but does not affect p65 DNA binding. **A**. Relative expression of genes implicated in invasion and metastasis, and that are known to be regulated via NF-κB activation in 4T1 spheroids. Fold change is calculated from normalized raw counts (RNA sequencing) of CPT1a knockout and control 4T1 3D spheroids cultured for 5 days in medium containing extra palmitate (75 μM) or acetate (5 mM). **B**. Relative expression of genes implicated in invasion and metastasis, and that are known to be regulated via NF-κB activation in MCF10A H-Ras^V12^ spheroids. Fold changes are calculated for CPT1a knockout and control MCF10A H-Ras^V12^ 3D spheroids cultured for 5 days in medium containing extra palmitate (75 μM) or acetate (5 mM) and are normalized to gene expression in control cells. One-way ANOVA with Dunnett’s multiple comparison test (n = 3). **C**. GSEA enrichment plots comparing the gene expression profiles in 4T1 3D spheroids transduced with a lentiviral vector containing sg*Cpt1a* or control sgGFP (top panel) and sg*Cpt1a* 4T1 3D spheroids cultured with or without acetate (bottom panel). NES, normalized enrichment score; the P value indicates the significance of the enrichment score. **D**. Total p65 binding to DNA measured by electrophoretic mobility shift assay (EMSA). Arrow indicates the position of the NF-κB containing complex. A representative of three experiments is shown.

**Supplemental Figure 6.**
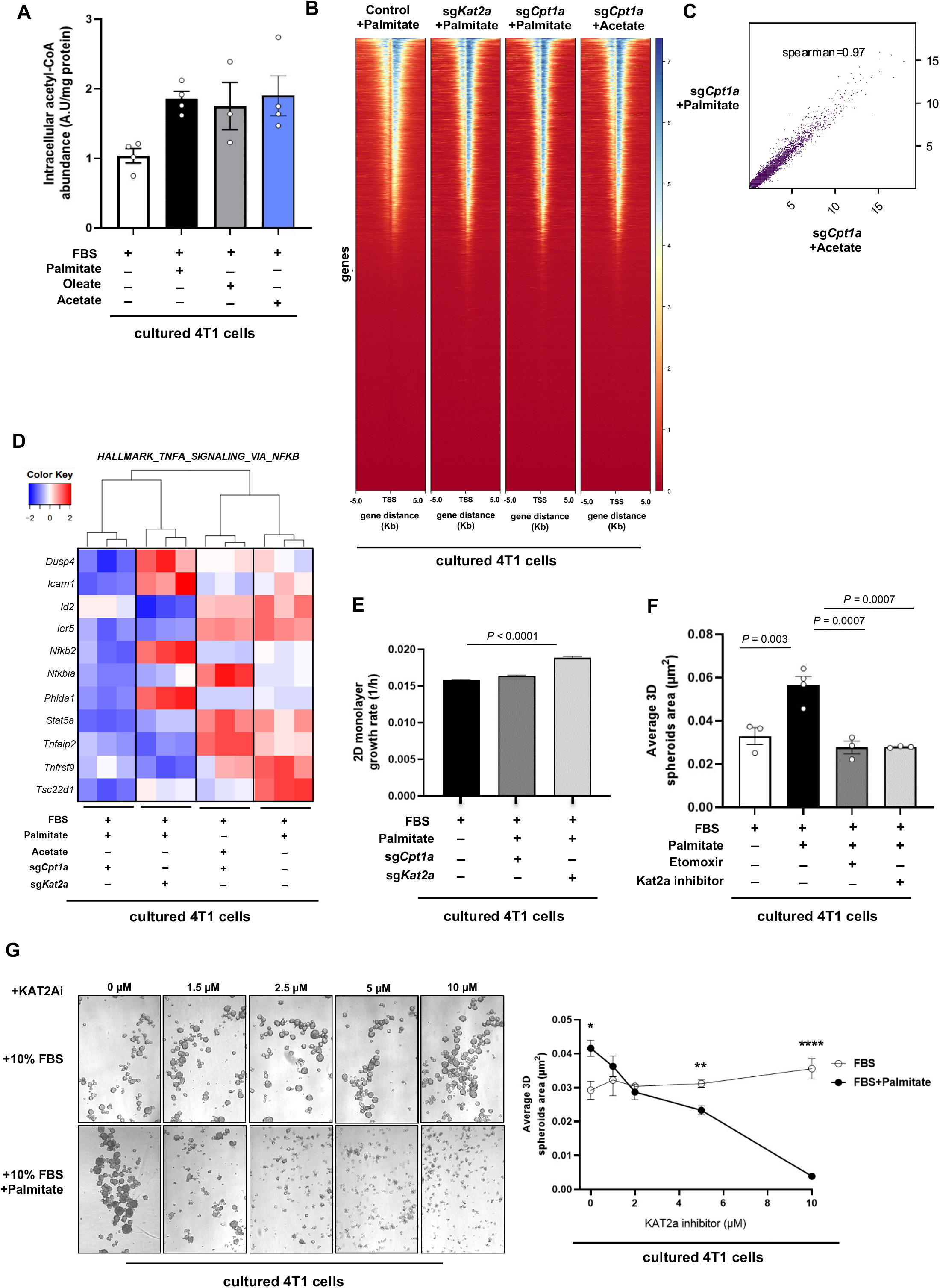
Global histone acetylation and chromatin accessibility are not consistently changed upon CPT1a and/or KAT2a loss. **A**. Intracellular levels of acetyl-CoA in 4T1 cells growing in 3D for 5 days in medium containing only 10% FBS, or supplemented with palmitate (75 Mm), oleate (116 μM) or acetate (5 mM). **B**. Heatmap of the signal intensity of H3K9ac-targeted gene loci in control, CPT1a and KAT2a knockout 4T1 3D spheroids cultured for 5 days in medium containing extra palmitate (75 μM). **C**. Correlation plot of H3K9 acetylation in 4T1 3D spheroids cultured in the presence of palmitate upon CPT1a inhibition with and without acetate. **D**. Heatmap and hierarchical clustering of top-scored downregulated genes of the NF-κB signaling pathway upon CPT1a deletion in 4T1 3D spheroids cultured in the presence of palmitate, represented together with the expression status of the same genes upon acetate rescue and KAT2a deletion. **E**. Growth rate in 4T1 cells upon genetic inhibition of either *Cpt1a* or *Kat2a* in 2D culture. One-way ANOVA with Dunnett’s multiple comparison test (n =6). **F**. 3D spheroids in 4T1 cells upon pharmacologic inhibition of either KAT2a using the inhibitor CPTH2 (2 μM) or CPT1A using etomoxir (50 μM) cultured for 5 days in medium with or without extra palmitate supplementation. Size quantification is represented by the average spheroids area of >100 spheroids. One-way ANOVA with Turkey’s multiple comparison test (n ≥4). **G**. Dose-response of 3D spheroid growth to the pharmacologic inhibition of KAT2a using CPTH2 inhibitor with or without extra supplementation of palmitate. *Left panel*, representative pictures. Right panel, spheroid size quantification is represented by the average spheroids area of >100 spheroids. Two-way ANOVA with Turkey’s multiple comparison test (n ≥4). Asterisks represent statistical significance as follows: *p < 0.05, **p < 0.01, ****p < 0.0001.

**Supplemental Figure 7.**
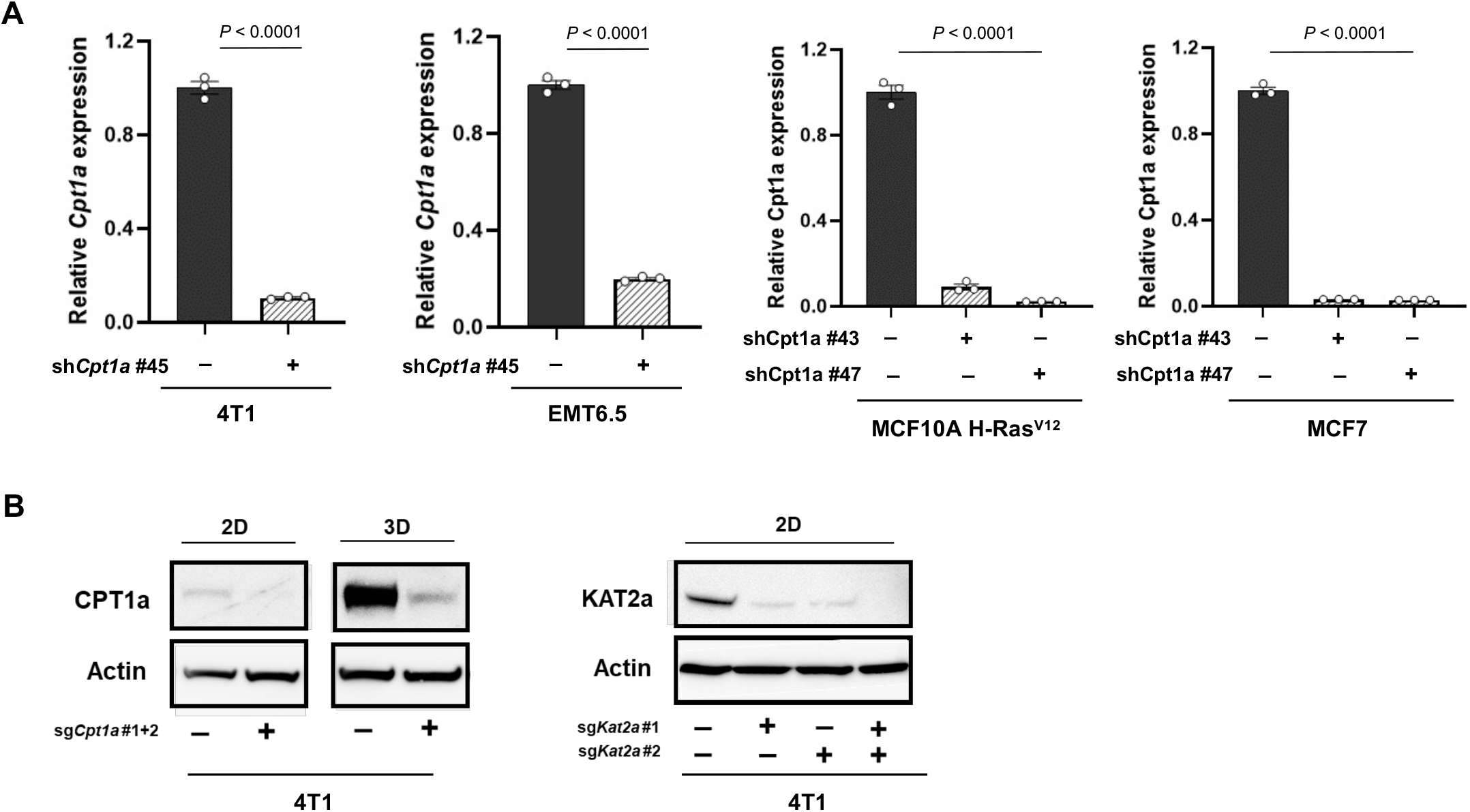
Protein and RNA expression of genetically modified breast cancer cells. **A**. Relative gene expression analysis of *CPT1A* in human (MCF10A H-Ras^V12^ and MCF7), mouse (4T1 and EMT6.5) breast cancer cells infected with either a control shRNA, or two different *CPT1A*, or*Cpt1a* shRNAs normalized to the control condition. **B**. Protein expression in mouse 4T1 cancer cells infected with either a control sgGFP or two different sgRNA against *Cpt1a* and *Kat2a* gRNAs.

## Material and methods

### Cell culture

MCF10A (ER-/PR-/HER2-), MCF7 (ER+/PR+/HER2-) and 4T1 (ER-/PR-/HER2-) cell lines were purchased from ATCC. The EMT6.5 (ER-/PR-/HER2-) cell line was provided by R. Anderson (Peter MacCallum Cancer Center). MCF10A cells expressing the oncogenic driver H-Ras^V12^ (MCF10A H-Ras^V12^) were generated as previously described^8^ as a relevant in vitro model for breast cancer since 50% of the human breast cancers display increased H-Ras activity ^75^. Moreover, the suitability of this cell line for metastasis biology has been previously reported ^8,9,76,77^. 4T1 and EMT6.5 cells were grown in Roswell Park Memorial Institute (RPMI) 1640 medium supplemented with 10% fetal bovine serum, 1% penicillin (50 units/mL and 1% streptomycin (50 μg/mL). MCF7 cells were maintained in Dulbecco’s modified Eagle’s medium (DMEM) supplemented with 10% fetal bovine serum (FBS), 1% penicillin (50 units/mL) and 1% streptomycin (50 μg/mL). MCF10A H-RAS^V12^ cells were cultured in DMEM/F12 supplemented with 5% horse serum, 1% penicillin (50 units/mL, 1% streptomycin (50 μg/mL), 0.5 μg/mL hydrocortisone, 100 ng/mL cholera toxin, 10 μg/mL insulin and 20 ng/mL recombinant human EGF. All cells were maintained at 37°C and 5% CO_2_ and 95% relative humidity and regularly tested negative for mycoplasma infection by Mycoalert detection kit (Lonza). For 3D growth conditions, plates covered with soft-agar were used as described^8^. Briefly, 1% soft-agar was mixed 1:1 with culture medium and left to solidify at room temperature. Cells were plated on top of the base agar and incubated for 3-5 days.

Sodium acetate was purchased from Sigma-Aldrich and used at a concentration of 5 mM. Sodium palmitate, stearic acid, oleic acid and octanoic acid were purchased from Sigma-Aldrich. These free fatty acids (FFA) were supplemented on top of FBS to the media in different concentrations up to complete a final concentration of 130 μM. FFA stock solutions were prepared by coupling free fatty acids with bovine serum albumin (BSA) as described previously^78^. The final ratio of FFA to BSA was always at least 3:1. Conditions with no additional FFA added were prepared using the same stocks of 10% w/w BSA with an equivalent amount of ethanol matching the concentration in FFA stocks. Etomoxir and the KAT2A specific inhibitor cyclopentylidene-[4-(4’-chlorophenyl)thiazol-2-yl)hydrazone (CPTH2)^79^ were purchased from Cayman Chemical, dissolved in ethanol and used at concentrations of 50 and 2 μM, respectively. The ATP Citrate lyase (ACLY) inhibitor BMS-303141 was purchased from MedChemExpress, dissolved in DMSO and used at 20 μM. Tumor Necrosis Factor-α from mouse was purchased from Sigma-Aldrich and used at 10 ng/μL. Ammonium pyrrolidinedithiocarbamate (PDTC) was purchased from Merck, dissolved in water and used at 0.5 μM.

### Cell proliferation assays

Growth was assessed based on cell number and cell confluency (two-dimensional, 2D) or spheroid size (three-dimensional, 3D). Specifically, to measure cell proliferation in 2D cultures, growth rates were calculated by measuring confluency every 4 hours using an Incucyte live-cell imager (Essen Biosciences, Ann Arbor, MI) over 96 hours in culture. To measure 3D growth, spheroids were cultured in 6-well plates upon specified conditions for 5 days. The compounds were always added at day 0 and representative pictures of each well were taken at day 5. Spheroids area was analyzed using Image Studio Lite 5.2 of ≥5 representative pictures per experimental condition. All growth experiments were performed in n≥3 biological replicates.

### Generation of shRNA knockdown cell lines

CPT1a knockdown cell lines were generated using the shRNA-expressing lentiviral pLKO-shRNA2 vector (No. PT4052-5; Clontech), with a puromycin selection cassette, kindly provided by Prof. P.Carmeliet, VIB, Belgium). Two non-overlapping oligonucleotide sequences per species (human and mouse) and nonsense scrambled shRNA sequence as negative control were used (oligonucleotide sequences are available upon request). Lentiviral particles were produced in HEK293 cells. Transduction of cells was performed overnight and the medium was replaced the next day. Polyclonal cells were selected for one week with puromycin, 1 μg/mL for MCF10A H-Ras^V12^, MCF7 and EMT6.5 and 2 μg/mL for 4T1 cells. shRNA-based silencing was confirmed by qPCR **(Supplemental Figure 7A**).

### Generation of CRISPR knockout cell lines

Murine *Cpt1a* and *Kat2a* knockout cell lines were generated by first establishing parental cell lines with doxycycline-inducible Cas9 expression using the Dharmacon Edit-R Inducible Lentiviral Cas9 vector (Horizon Discovery). Complementary sgRNA oligonucleotides targeting two different exons of either *Cpt1a* or *Kat2a* were designed and cloned into the LentiGuide Puro sgRNA expression vector (Addgene plasmid #52963) and delivered to cells via lentiviral infection. Targeting sequences - mouse Cpt1a sgRNA1 (CACATTGTCGTGTACCACAG), mouse Cpt1a sgRNA2 (ACGTTGGACGAATCGGAACA), mouse Kat2a sgRNA1 (GCTTCGGCCAAACACGTGGG), mouse Kat2a sgRNA2 (GCTCGCCTGGAAGAACGGCG), and non-targeting sgRNA against EGFP (GAGCTGGACGGCGACGTAAA) - were synthesized by IDT. Lentiviral particles were produced in HEK293T cells. Stable 4T1 cell lines were generated by selection with puromycin (2 μg/mL). Cas9 expression was induced using doxycycline (2ug/ml) for 7 days after which gene knockout was verified by western blot **(Supplemental Figure 7B)**.

### Gene expression analysis by quantitative PCR

Total RNA from frozen tissues or cell lines was isolated using Trizol and cDNA synthesis was performed using qScript cDNA Synthesis Kit (Quanta) according to the manufacturer’s protocols. 50 ng of cDNA per reaction were amplified at 95⍰°C for 10 min, followed by 40 cycles of 15 s at 95⍰°C and 1 min at 60⍰°C using SYBER Green PCR Master Mix (Life Technologies) in the 7500HT system (Applied Biosystems). The mRNA expression in each sample was normalized to endogenous housekeeping gene *RPL19* and relative expression was analyzed using 2^−ΔΔCt^ method. Gene-specific primers were designed using Primer3 or obtained from Origene. Sequences are listed in Table 1.

**Table 1.**
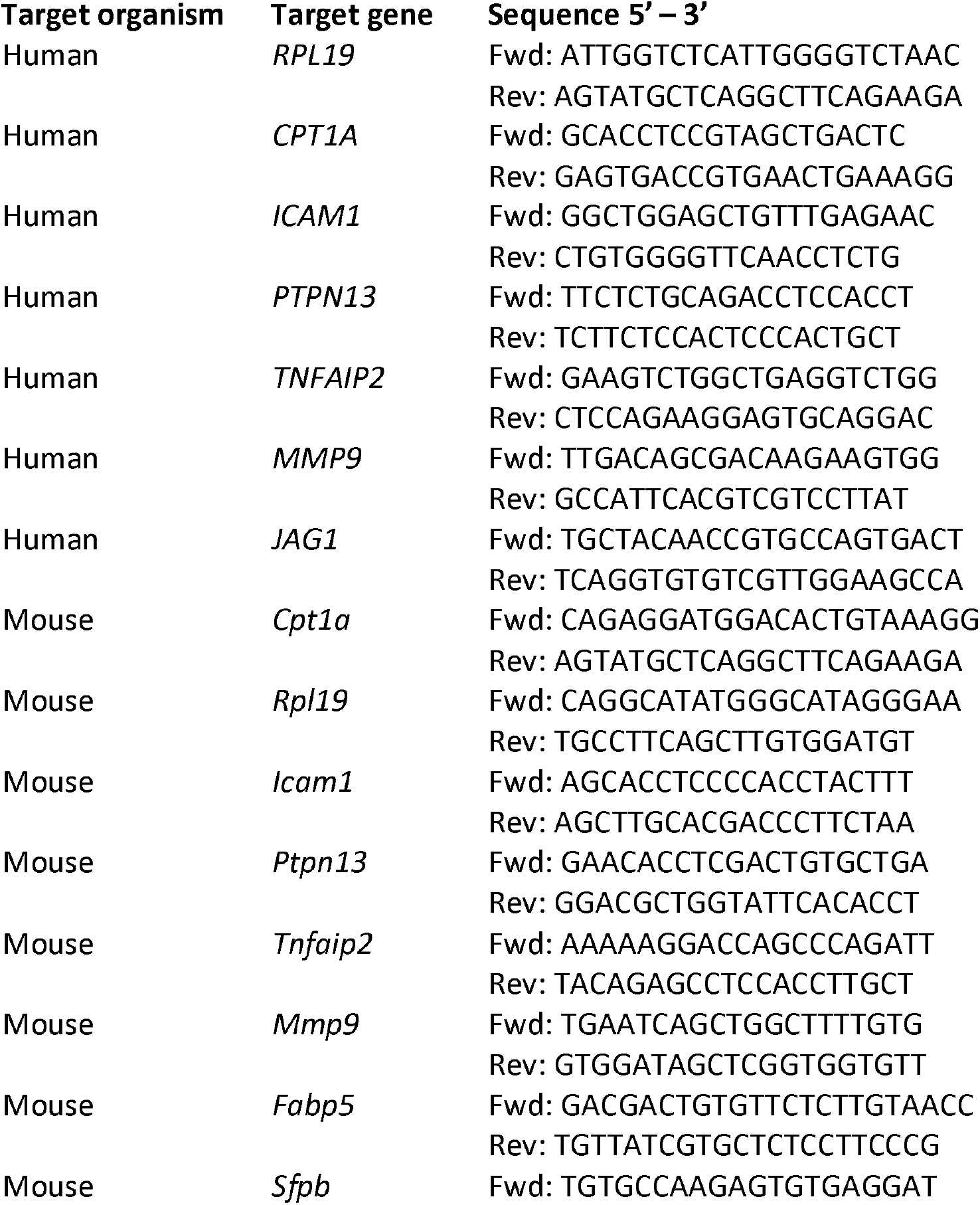
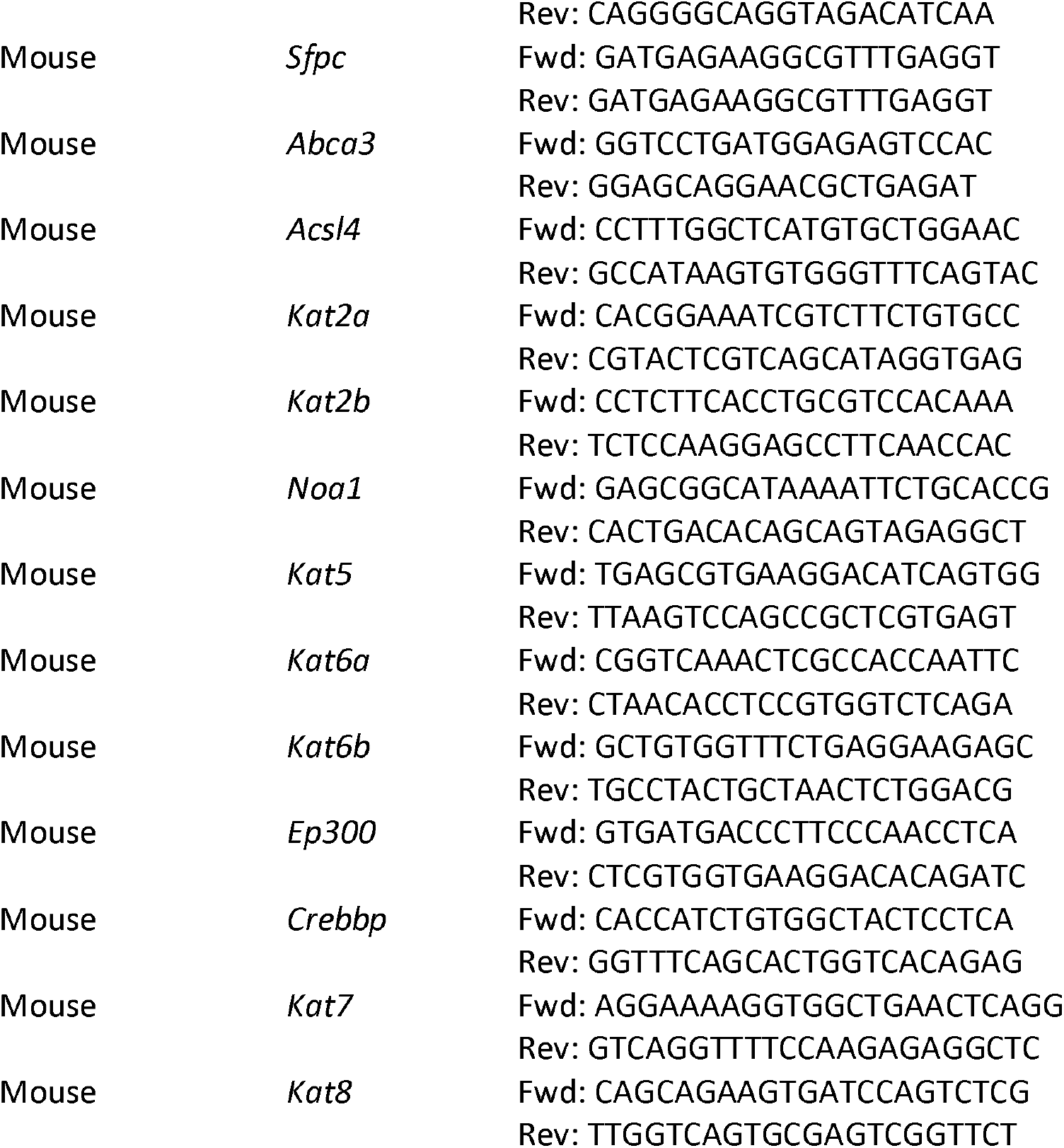
Primers used to analyze mRNA levels

### Cell-fractionation and nuclear isolation

Nuclear and cytoplasmic extracts were prepared from spheroids as described previously^80^. Briefly, spheroids were collected and washed once in ice-cold PBS. Spheroid pellets were then suspended in 1.5 ml of hypotonic cytoplasmic fraction buffer (20 mM HEPES pH 8.0, 0.2 % IGEPAL CA-630, 1 mM EDTA, 1 mM DTT, protease and phosphatase inhibitor cocktail), lyse using tissue-lyser and incubated on ice for 10 min with occasional shaking. After centrifugation (4 °C, 1700⍰× ⍰g, 5 min), supernatant was set aside and treated as the cytoplasmic fraction. The remaining pellet was suspended in 100 μl of nuclear fraction buffer (20 mM HEPES pH 8, 420 mM NaCl, 20 % glycerol, 1 mM EDTA, 1 mM DTT, protein and phosphatase inhibitors), incubated on ice for 30 min with occasional mixing and then centrifuged at 4 °C, 10,00⍰× ⍰g, 10 min. Supernatant was transferred to a new tube and treated as nuclear fraction. Protein concentration was measured using Bradford Assay.

### Protein extraction and western blot analysis

Spheroids were collected by centrifugation and lysed in RIPA lysis buffer (Thermo Scientific) supplemented with protease (complete, Mini Protease Inhibitor Cocktail Tablets (Roche) and phosphatase (PhosSTOP™, Sigma) inhibitors. Tissues were ground prior RIPA incubation. Protein amount was measured using a Pierce BCA protein assay kit (Thermo Scientific). Aliquots of 30-40⍰μg of protein were loaded on a NuPAGE 4–12% denaturing Bis-Tris gel and transferred to a nitrocellulose membrane (Thermo Scientific). Membranes were incubated for 1h at room temperature in a blocking solution of 5% milk in TRIS Buffer Saline 0.05% Tween (TBS-T). Subsequently, membranes were incubated overnight at 4°C with the following primary antibodies: CPT1a (Abcam, ab234111; 1:1000 dilution), GCN5L2/KAT2A (C26A10) (Cell signaling, 3305T; 1:1000 dilution), NF-kB p65 (acetyl K310) (Abcam, ab19870; 1:1000 dilution), NF-κB p65 (D14E12) (Cell signaling, 8242S; 1:1000 dilution), β-actin (Sigma, A5441; 1:10,000 dilution) or Histone H3 (Abcam, ab1791; 1:2000 dilution). The day after the membranes were incubated with a mouse (Cell Signaling Technology, 7076; 1:4000 dilution) or rabbit (Cell Signaling Technology, 7076; 1:4000 dilution) HRP-linked secondary antibodies, and bound antibodies were visualized using Pierce ECL reagent (Thermo Scientific). Images were acquired using an ImageQuant LAS 4000 (GE Healthcare).

### Electrophoretic Mobility Shift Assays (EMSA)

Whole extracts from 3D spheroids were prepared and analyzed for DNA binding activity using the HIV-LTR tandem κB oligonucleotide as κB probe as described previously^81^. Use of whole cell extracts to analyze NF-κB DNA-binding status by EMSA was previously validated by comparing nuclear and total NF-κB DNA-binding profiles and the observation of highly similar NF-κB DNA-binding activity, indicating that NF-κB binding emanates from nuclear proteins ^82^.

### Mass spectrometry analysis

To analyze metabolite abundances of three-dimensional (3D) spheroids, quenching and extraction procedures were applied as described previously ^83^. Briefly, spheroids growing in 6-well plates were collected after 5 days by centrifugation (1.5 min at 1400 rpm). Pellets were quenched by transferring them to a tube placed inside of a rack in a cold ethanol bath (–40°C) that contained cold quenching solution (60% methanol, 10% ammonium acetate (100 mM) and 30% Milli-Q). Spheroids were spun down (1 min at 2500 rpm), washed using −20°C cold quenching solution and placed in dry ice until continuing with the extraction. To analyze two-dimensional (2D) cultured cells, cells were seeded in 6-well plates and, after 5 days, cells were washed with blood bank saline and quenched by flash-freezing the plates in liquid nitrogen. Metabolites for mass spectrometry analysis were extracted, derivatized and measured as described previously ^84,85^. Briefly, for 2D cells, 800 μL of −20 °C cold 62.5% methanol-water buffer containing glutarate as internal standard (2.5 μg/mL) was added to the quenched wells and cells were scraped using a pipette tip. Suspensions were transferred to Eppendorf tubes and 500 μL of −20°C cold chloroform containing heptadecanoate (10 μg/mL) as internal standard was added. For 3D spheroids, 800 μL of −20 °C cold 62.5% methanol-water buffer was added into Eppendorfs containing the quenched spheroids and a tissue-lyser was applied inside of the cold ethanol bath (–40°C) until the spheroids were completely disintegrated. After, 500 μL of −20°C cold chloroform was added. 2D and 3D samples were vortexed at 4°C for 10 min to extract metabolites. Phase separation was achieved by centrifugation at 4°C for 10 min max rpm, after which the chloroform phase (containing the total fatty acid content) and the methanol phase (containing polar metabolites) were separated and dried by vacuum centrifugation.

Metabolite abundances were analyzed by either gas or liquid chromatography-mass spectrometry. For fatty acid measurements, the lipid fraction (chloroform phase) was esterified with 500 μL 2% sulfuric acid in methanol at 60°C overnight and extracted by addition of 600 μL hexane and 100 μL saturated aqueous NaCl. Samples were centrifuged for 5 min and the hexane phase was separated and dried by vacuum centrifugation. Samples were resuspended in hexane and fatty acids were separated with gas chromatography (8860 or 7890A GC system, Agilent Technologies, CA, USA) combined with mass spectrometry (5977B or 5975C Inert MS system, Agilent Technologies, CA, USA). 1 μL of each sample was injected in splitless mode, split ratio 1 to 3 or 1 to 9 with an inlet temperature of 270 °C on a DB-FASTFAME column (30m x 0.250 mm). Helium was used as a carrier gas with a flow rate of 1 mL per min. For the separation of fatty acids, the initial gradient temperature was set at 50°C for 1 min and increased at the ramping rate of 12°C/min to 180°C, followed by a ramping rate of 1°C/min to rich 200°C for 1 min. The final gradient temperature was set at 230°C with a ramping rate of 5°C/min for 2 min. The temperatures of the quadrupole and the source were set at 150°C and 230°C, respectively. The mass spectrometry system was operated under electron impact ionization at 70 eV and a mass range of 100-600 amu was scanned. For acetyl-CoA and polar metabolites measurements, the polar metabolite phase was resuspended in 50 μL water and analyzed in a Dionex UltiMate 3000 LC System (Thermo Scientific) with a thermal autosampler set at 4°C, coupled to a Q Exactive Orbitrap mass spectrometer (Thermo Scientific) and a volume of 10 μL of sample was injected on a C18 column (Acquity UPLC HSS T3 1.8μm 2.1 × 100mm). The separation of metabolites was achieved at 40°C with a flow rate of 0.25 ml/min. A gradient was applied for 40 min (solvent A: 10mM Tributyl-Amine, 15mM acetic acid – solvent B: Methanol) to separate the targeted metabolites (0 min: 0% B, 2 min: 0% B, 7 min: 37% B, 14 min: 41% B, 26 min: 100% B, 30 min: 100% B, 31 min: 0% B; 40 min: 0% B. The MS operated in negative full scan mode (m/z range: 70-500 and 190-300 from 5 to 25 min) using a spray voltage of 4.9 kV, capillary temperature of 320°C, sheath gas at 50.0, auxiliary gas at 10.0. Data was collected using the Xcalibur software (Thermo Scientific). For the detection of metabolites by LC-MS/MS, a 1290 Infinity II with a thermal autosampler set at 4°C, coupled to a 6470 triple quadrupole (Agilent Technologies) was used. Samples were resuspended in 80% methanol-water and a volume of 4 μL was injected on a SeQuant ZIC/pHILIC Polymeric column (Merck Millipore). The separation of metabolites was achieved at 25°C with a flow rate of 0.20 ml/min. A gradient was applied for 22 min (solvent A: 10 mM ammonium acetate (pH=9.3, 10╰mM) – solvent B: acetonitrile) to separate the targeted metabolites (0 min: 10% A, 2 min: 10% A, 13 min: 30% A, 13.1 min: 70% A, 17 min: 75% A, 18 min: 10% A, 22 min: 10% A).The temperature of the gas and the sheath gas were set at 270°C (flow: 10L/min) and 300°C (flow: 12L/min), respectively.

Metabolite abundances were calculated from the raw chromatograms using an in-house MATLAB script. For relative abundance, total ion counts were normalized to an internal standard (glutarate or heptadecanoate) and the protein content for cell extracts. To calculate fatty acid concentration in interstitial fluid, a known volume of interstitial fluid was extracted using 62.5% methanol-water and chloroform buffers as described above. A standard curve for each metabolite was extracted and analyzed in parallel.

### Histone extraction and SILAC analysis

For the preparation of the SILAC labeled histone internal standard, 4T1 cells were cultured in SILAC DMEM (Silantes) supplemented with 10% 10 kDa dialyzed FBS (PAA), 2 mM glutamine, 1% penicillin/streptomycin, 84 mg/l 13C615N4 L-arginine and 175 mg/l 13C615N2 L-lysine (Cambridge Isotope Laboratories) until they were fully labeled. 3D spheroids were collected after 5 days and the core histone proteins were isolated by acid extraction procedure previously described (Karch et al. 2016). Briefly, nuclei were isolated from pelleted cells with a high salt extraction buffer (15 mM Tris-HCL (pH 7.5), 60 mM KCl, 15 mM NaCl, 5 mM MgCl2, 1 mM CaCl2, 250 mM Sucrose, 0.3% NP-40). Histones were isolated from nuclei by incubation in 0.4 N H2SO4, followed by addition of trichloroacetic acid to a final concentration of 20%. Isolated histones were precipitated with acetone and resuspended in 8 M urea. SILAC labeled histones were added to unlabelled histones as an internal standard in a 2:1 ratio. Histones were digested in 50 mM ammonium bicarbonate using Arg-C (Promega), and the resulting peptides desalted by C18 StageTip ^86^. Peptides were resuspended in 1% TFA, 0.2% formic acid buffer and injected on an EASY-nLC (Thermo Fisher Scientific) coupled online to a mass spectrometer. Peptides were separated on a 20 cm fused silica emitter (New Objective) packed in-house with reverse-phase Reprosil Pur Basic 1.9 μm (Dr. Maisch GmbH). Peptides were eluted with a flow of 300 nl/min from 5% to 30% of buffer B (80% acetonitrile, 0.1% formic acid) in a 60 min linear gradient. Eluted peptides were injected into an Orbitrap Fusion Lumos (Thermo Fisher Scientific) via electrospray ionisation. MS data were acquired using XCalibur software (Thermo Fisher Scientific). The MS .raw files were processed with MaxQuant software (version 1.6.6.3) and searched with the Andromeda search engine with the following settings: minimal peptide length 6 amino acids, fixed modification Carbamidomethyl (C) and variable modifications Acetyl (K), Acetyl (Protein N-term) and Oxidation (M). Multiplicity was set to 2, where the light labels were Arg0 and Lys0 and the heavy labels were Arg10 and Lys8. The false discovery rates (FDRs) at the protein and peptide level were set to 1%. Perseus (version 1.6.2.2) was used for downstream analysis. The data were filtered to remove potential contaminants, reverse peptides which match a decoy database, and proteins only identified by site. To ensure unambiguous identification, only proteins identified with at least one unique peptide were considered. The heavy/light SILAC ratios were transformed by 1/x and then by log2. Each acetylation site was normalized by total histone abundance.

### ChIP sequencing analysis

To analyze changes in genome-wide DNA binding sites using, we applied ChIPmentation protocol followed by sequencing as described previously (Broux et al. 2019). Briefly, 3D spheroids were collected after 5 days, cross-linked with 1% formaldehyde for 10 min at room temperature and then quenched by addition of glycine (125 mM final concentration). Nuclei were isolated and resuspended in L3B+ buffer (10 mM Tris-Cl pH 8.0, 100 mM NaCl, 1 mM EDTA, 0.5 mM EGTA, 0.1% sodium deoxycholate, 0.5% N-Lauroylsarcosine, 0.2% SDS). Chromatin was fragmented to 100-300 bp using 25 cycles (30 sec on, 30 sec off, 20% amplitude) using Qsonica Q800R. Chromatin immunoprecipitation was performed overnight in the presence of H3K9ac antibody (Cell signaling, 5685S) conjugated to magnetic protein A/G beads (Millipore). Drosophila melanogaster chromatin was spiked-in (50 ng, Active Motif, 53083). Next, tagmentation (Nextera DNA Sample Preparation Kit, Illumina) and library preparation was performed, then DNA was purified using three sided SPRI bead cleanup using 1.0X; 0.65X; 0.9X ratios (Agencourt AMPure Beads, Beckman Coulter) and sequenced on Illumina Hiseq 4000 platform (Illumina, San Diego, CA, USA). The raw sequencing reads were cleaned with fastq-mcf and a quality control was performed with FastQC. These cleaned reads were then aligned to the mm10 genome using BWA and duplicates were removed with Picard. deepTools was used to plot heatmaps of signals centered around TSS as well as to plot Spearman correlation of ChIP-seq signal per gene between conditions.

### In vivo experiments using BALB/c mice

All animal experiments complied with ethical regulations and were approved by the Institutional Animal Care and Research Advisory Committee of KU Leuven, Belgium (ECD number P007/2020 and ECD P025/2020). Sample size was determined using power calculations with B = 0.8 and P < 0.05 based on preliminary data and in compliance with the 3R system: Replacement, Reduction, Refinement. 6-8 weeks old female BALB/c mice were inoculated with 4T1 or EMT6.5 cells either in the mammary fat pad (m.f.; 1 × 10^6^ cells) or intravenously (i.v.; 1 × 10^5^ cells). For etomoxir treatment, 1×10^6^ 4T1 cells were suspended in 50 μL of PBS and injected in the mammary fat pad of BALB/c mice. Mice were checked for tumor formation by palpation, and when tumors were about 13 mm, mice were randomized into two groups for intraperitoneal injections daily for 3 days: vehicle (water) and treatment (etomoxir, 40 mg/kg per day). Mice were sacrificed and lung metastases were collected to further analysis. For metabolomics, mice were euthanized using Nembutal, and tumor tissues were placed in cold saline for rapid dissection (>3 min) and immediately frozen using a liquid-nitrogen-cooled Biosqueezer (Biospec Products). The tissue was weighed (10–15 mg) and pulverized (Cryomill, Retsch) under liquid-nitrogen conditions. To follow the effect on the primary tumor, tumor volumes were measured during the experiment using a caliper. Additionally, at the end of the experiment, primary tumors were dissected and weighted. Humane endpoints were determined as follows: tumor size of 1.8 cm^3^, loss of ability to ambulate, labored respiration, surgical infection, or weight loss over 10% of initial body weight. Mice were monitored and upon detection of one of the previously mentioned symptoms, the animal was euthanized.

For high fat diet experiments, 4-weeks old female BALB/c mice were randomized into two groups: control diet (CD, E15742-33 ssniff Spezialdiäten GmbH) or high fat diet (HFD, S8655-E220 sniff Spezialdiäten GmbH). The energy balance between fat/protein/carbohydrates was 13%/27%/60% and 60%/20%/20% for CD and HFD, respectively. Mice were maintained in these diets for 16 weeks. Bodyweight was monitored biweekly and mice were inspected weekly for welfare assessment.

### Lung and liver interstitial fluid extraction

Interstitial fluid extraction method was adapted and performed as described previously^7,22^. For the collection of human samples, ‘normal’ lung tissue was collected from patients who underwent lung surgery for emphysematous lung volume reduction or tumorectomy (taken as far away as possible from the tumor front). The study was approved by the local ethics committee (Medical Ethics Committee UZ/KU Leuven) under the protocol S57123. For the collection of mice samples, 6-8 balb/c female mice were first euthanatized with 50μL of 60mg/ml of Dolethal (pentobarbital sodium), and subsequently, liver and lung were harvested by surgical resection, washed with a blood bank saline solution and dried from liquid excess by carefully tapping in a gauzeSubsequently, the organs were placed in a homemade filtered centrifuge tube with a nylon mesh filter with 20 μm opening pores (Repligen). The interstitial fluid (between 1-5 μL for each organ) was collected in the column-centrifuge tubes by centrifugation at 400g, 4°C for 10 minutes and stored in dry ice immediately after extraction. The volume of interstitial fluid was used to determine the concentration of the different metabolites measured by mass spectrometry.

### Tumor condition media generation and in vivo pre-metastatic niche formation

The procedure to perform pre-metastatic niche formation was adapted from ^25^. In brief, for primary tumor formation, 1×10^6^ 4T1 cells were injected orthotopically in the mammary fat pad of BALB/c mice (7-8 weeks old). After 21 days, mice were euthanized and primary tumor was removed. Tumors were cut in small pieces and placed in 10-cm dishes cultured with 15 mL of DMEM/F12 + 1% penicillin (50 units/mL) and 1% streptomycin (50 μg/mL). Tumors were incubated 72h at 37°C and 5% CO_2_. The same media was incubated in parallel in 10cm dishes without tumor to generate control media. After 72h, tumor- and control-conditioned media were collected, transferred to a cell strainer (70 μm) and spin down 10 min at 4000 rpm. Supernatants were moved to 50 mL tubes by pooling together conditioned media of the same type from 3 different tumors to cover the variability between tumors. Hepes (20mM) was added to the media and this was filtered (0.20 μm) and aliquoted to be stored at 4*C no more than 2 weeks after collection. 7-8 weeks old balb/c mice were injected intravenously with 200 uL of either control or tumor conditioned media 3 times per week for 3 weeks. Three days after the last injections, interstitial fluid of the lung was collected as described previously.

### Haematoxylin and eosin staining of tumor sections

Haematoxylin and eosin (H&E) staining of pulmonary metastases was performed as previously described^9^. Briefly, dissected lung samples were gently infused via the trachea with 10% formalin, fixed overnight, and immersed in 70% ethanol for 24 hours. Then, 5-μm thick sections obtained from the resulting paraffin blocks were stained with haematoxylin and eosin. Scanned slides were analyzed for metastatic area and the number of metastases using ZEN blue software. Metastatic index was calculated by dividing the metastatic area by the primary tumor weight. Only mice with a primary tumor of 0.8 g or greater were analyzed. All i.v.-injected animals were analyzed. All samples were analyzed blinded.

### Lung dissociation and flow cytometry analysis of cancer cell populations

Prior injections, 4T1 and EMT6.5 cancer cells were transfected with the mammalian expression lentiviral vector pLKO.3 Thy1.1 (Addgene plasmid #14749) containing the surface protein Thy1.1 as a reporter protein. Since Thy1.1 is not usually expressed on mouse cells and does not elicit immune activation if injected in recipient mice, its overexpression served to identify cancer cells from single-cell suspension of mouse tissues. Lentiviral particles were produced in HEK293T cells and transduction of cancer cells was performed overnight. After 72 hours of recovering, CD90.1-positive cells were FACS-sorted. CD90.1-expressing 4T1 and EMT6.5 cells were then injected i.v into 6-8 weeks old BALB/c mice. After 11 days, mice were anesthetized with Ketamine (100mg/kg) + Xylazne (10mg/kg) and lungs were perfused through the right ventricle (adapted from ^87^). Once lungs were extracted, the tissue was washed in blood bank saline, dry and minced for >2min using blades. Tissues were incubated with Liberase (Roche) (0.3 mg/mL) and DNAse1 (1 μg/mL) during 45 min at 37°C with occasional vortex. The reaction was quenched with 3% FBS:PBS + 2mM EDTA and filtered through a 70μm cell strainer. Cell pellet was washed, incubated with Red Blood Lysis buffer (Merck) and transferred through a 40 μm cell strainer. Single-cells suspension was counted and 30×10^6^ cells/mL were stained for flow cytometry analysis. Antibodies against CD45 (BD Bioscience, 550994 1:250 dilution), PDPN (BioLegend, 127409 1:250 dilution) and CD90.1 (BioLegend, 202505 1:400 dilution) were used for selecting cancer cells from the single-cell suspensions of murine lung. For cell-surface antigen staining, the samples were incubated with antibodies for 20 min at 4°C, washed and resuspended in PBS containing 3% FBS:PBS +2 mM EDTA. To exclude dead cells, Viability efluor450 (ThermoFisher, 65-0863-14 1:500 dilution) was used. Single cells were analyzed using a BD FACSCanto II with FACSDiva (BD Biosciences).

### Primary alveolar type II cells isolation by flow cytometry and 3D culture

Primary AT2 cells were isolated from CreER induced lineage-tracing mice^88^. Sftpc-CreERT2 mice were a gift from the J-H Lee laboratory (MRC Cambridge Stem Cell Institute). Breeding and all animal procedures at the Francis Crick Institute were performed in accordance with UK Home Office regulations under project license PPL80/2531. For type II cell lineage labeling, 3 doses of tamoxifen dissolved in Mazola corn oil (40 mg/mL stock solution) were given to *Sftpc-CreERT2;R26R-YFP* female C57BL/6J mice (8-10 weeks) via oral gavage over consecutive days (0.2 mg per g body weight). Lungs were harvested two weeks after the final dose. Lung tissues were dissociated as previously described ^37^. Briefly, lungs were minced manually and then digested in a mixture of DNase I (Merck Sigma-Aldrich) and Liberase TM and TH (Roche) in HBSS solution for 30 min in a 180rpm shaker at 37ºC. Samples were then washed, passed through a 100 μm filter, and centrifuged at 1250 rpm for 10 min. The resulting cell pellet was incubated in Red Blood Cell Lysis buffer (Miltenyi Biotec) for 4-5 min at room temperature. After a wash with MACS buffer (0.5% BSA and 250 mM EDTA in PBS), samples were passed through a 20μm strainer-capped flow cytometry tube to generate a single cell suspension. Cells were incubated with mouse FcR Blocking Reagent (Miltenyi Biotec) for 10 min at 4°C followed by an incubation with CD45-BV421 (Biolegend 103133, clone 30-F11, 1:200 dilution) and Ter119-BV421 (Biolegend 116233, TER-119, 1:200 dilution) for 30 min at 4ºC. Cells were washed twice with MACS buffer, before being incubated with 4⍰,6-diamidino-2-phenylindole (DAPI) to discriminate dead cells. AT2 cells (CD45-Ter119-YFP+) were sorted on a BD Influx cell sorter (BD Biosciences).

Isolated AT2 cells were seeded at a density of 150,000 cells/well in a collagen-coated Alvetex™ Scaffold 12-well plate insert (ReproCELL). Collagen solution was made by 30μg/mL PureCol collagen (Advanced Biomatrix), 0.1% bovine serum albumin, 20mM HEPES in HBSS. 2 mL of control or tumour-conditioned (generated as described above) MEM medium (DMEM/F12) with 0.5% fetal bovine serum, 100 U/ml penicillin-streptomycin, 20 ng/ml EGF and 10 μg/ml insulin was added below the insert. Cells were cultured at 37ºC and 5% CO_2_ for 72h. Media was collected at experimental endpoint, centrifuged and filtered to remove any cellular debris, and stored at -80°C until further metabolomics analysis by mass spectrometry. For RNA extraction, scaffolds were washed and placed directly into QIAzol lysis reagent and incubated on a shaker at 100 rpm for 15 minutes at room temperature. RNA was then isolated using the RNeasy Mini kit (Qiagen), together with an on-column DNase treatment (Qiagen). cDNA was synthesized with random primers, using the SuperScript III First Strand Synthesis Kit (Thermo Fisher Scientific) according to the manufacturer’s protocol.

### Single-cell RNA sequencing, data pre-processing, and cell-type assignment

Single cell suspensions from lungs of BALB/c mice injected with control media or tumor conditioned media for 3 weeks were achieved as described following the protocol described for lung dissociation. Cell suspensions were kept on ice and immediately processed for single-cell library preparation. Cell suspensions for each sample were converted to barcoded single-cell cDNA libraries using the Chromium Single Cell 5’ V1.1 Library (10x Genomics), following the manufacturer’s guidelines, and aiming for a total of 10000 cells per library. Single-cell libraries were then sequenced on a NovaSeq 6000 System (Illumina). The sequenced reads were then mapped to the mouse genome (mm10 build GRCm38.p6) using the Cell Ranger software (10x Genomics), and the resulting single-cell gene expression data were analyzed within the *R/Bioconductor* framework. Specifically, the raw UMI count matrices for each individual sample were first imported using *Seurat* ^89^, and then converted for further processing with *Monocle* ^90^. Low-quality cells were then filtered based on standard quality-control metrics, with sample-specific thresholds chosen based on evaluating quality-control histograms for each sample independently. In particular, cells were filtered based on their mitochondrial RNA content (allowing for a maximum of 10% in all cases), library size (removing cells with total UMI counts below 500 in both cases and number of detected genes (removing cells expressing less than 200 genes in both cases. Genes expressed in less than 5 cells were additionally ignored in all subsequent analyses. Size-factor and variance-stabilizing normalization (based on fitting to a negative binomial distribution) were then applied to the filtered data sets, and highly variable genes (HVGs) were identified for each of them based on their departure from the average normalized dispersion *versus* expression trend observed among all genes. After excluding mitochondrial, ribosomal-protein, and cell cycle-associated genes, the top 1000 HVGs with size-factor normalized expressions above 0.0035 (CM) and 0.001 (TCM) were selected. Principal component analysis (PCA) was then performed on the size factor-normalized and variance-stabilized count matrix restricted to these genes only, followed by 2D UMAP dimensional reduction ^90^ based on the resulting top 50 principal components (with *correlation distance metric, number of neighbors* = 15, and *minimum* distance = 0.1, and without further PCA scaling). After that, cells were clustered in the UMAP plane by applying the *Louvain* ^90,91^ graph-based algorithm at high resolution (resolution = 0.001 with k_NN_ = 8/7 for the CM/TCM samples, respectively), in order to attain a fine-grained cluster structure for each sample (90/62 clusters for the CM and TCM samples, respectively). The resulting fine-grained clusters were then manually annotated to specific cell types, based on evaluating the cluster-averaged normalized expression profiles of several cell type-specific markers. For marker score analysis, cell-type marker scores were calculated using the GSVA (gene set variation analysis) package ^92^. In brief, count matrices for the filtered data sets were first subject to size-factor and variance-stabilizing normalization. GSVA-based z-scores were then determined for a manually assembled list of marker gene sets). Scores for each marker set were then determined for every cell by normalizing the GSVA z-scores to the range 0–1, by means of the following mapping:

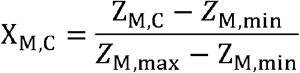

where 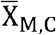 is the scaled (0–1) score for a given marker set, M in the cell C, while Z_M,C_ is the az-score for that cell, and Z_M,max_ and Z_M,min_ are respectively the highest and lowest values of z_M,C_ among all cells.

### Patient selection and sample collection

This study has benefited from two different clinical programs developed at the UZ Leuven hospital (Belgium).

#### UPTIDER program

Snap frozen and/or fresh tissue samples from metastases as well as normal tissues were obtained through the ethically approved UPTIDER program (UZ/KU Leuven Program for Post-mortem Tissue Donation to Enhance Research, NCT04531696, S64410). In this project, patients with metastatic breast cancer that consent to participate undergo a rapid research autopsy in the first 12 hours after death. In short, upon death of a participating patient, the body is transported to the morgue of the UZ Leuven hospital. Several types of body fluids are collected, as well as extensive malignant and adjacent normal tissue samples from different organs in the body, including the ones that are difficult to reach during the life of the patient. Tissue samples are processed in different conditions (snap-frozen in liquid nitrogen, frozen in OCT (Optimal Cutting Temperature Compound), collected in neutral buffered formalin for further processing in formalin-fixed paraffin embedded blocks (FFPE), or fresh in specific media for in vivo development). For our experiments described in this paper, fresh samples were collected and others snap-frozen in liquid nitrogen. Clinicopathological information for patients 2004 and 2009 in the UPTIDER data set is shown in Supplemental Table 1. The UPTIDER project is coordinated and performed by the Laboratory for Translational Breast Cancer Research (LTBCR) at our institution, KU Leuven, under the lead of Prof. Christine Desmedt and Prof.

Dr. Giuseppe Floris. The study was approved by the KU-UZ Leuven ethical committee and informed consent was obtained from patients.

#### CHEMOREL program

Patients for the retrospective translational CHEMOREL study were selected from a clinical-pathological database in which all breast cancer patients, who were diagnosed and treated at the UZ Leuven Multidisciplinary Breast Center, are documented (i.e. patient and tumor characteristics, therapy, and follow-up information including relapse and survival). From this database, a homogeneous cohort of newly diagnosed primary Luminal B-type breast cancer patients with grade 2/3, ER+, PR+/-, HER2-tumors was selected including (i) patients who remained disease-free for at least 6 to 10 years after the initial therapy (n=43), (ii) patients developing distant metastasis within 5 years after initial therapy (n=44), and (iii) primary metastasized patients (n=14). Groups were matched for patient age, tumor size, and node status. Clinicopathological information for patients in the CHEMOREL data set is shown in Supplemental Table 2. The study was approved by the KU-UZ Leuven ethical committee and informed consent was obtained from patients.

### RNA sequencing analysis

#### Patient tumors

RNA was extracted from 5×10μm FFPE unstained tissue slides prepared from the left-over of surgical resection specimens after standard pathological diagnostic procedures. An extra tissue slide was cut sequentially with the unstained slides and was H&E stained and microscopically inspected by an expert breast pathologist (Prof. Dr. G. Floris, UZ Leuven) to ensure representative and comparable tumor cellular composition across the entire cohort. RNA extraction was performed by using the HighPure FFPET RNA extraction kit (Roche). RNA concentration and quality were assessed on the Agilent Bioanalyzer. RNA sequencing workflow and subsequent bioinformatics were accomplished at the Laboratory of Translational Genetics (VIB-KU Leuven, Prof. D. Lambrechts). RNA libraries were created using the KAPA stranded mRNA seq Library preparation kit according to the manufacturer’s instructions. Briefly, poly-A containing mRNA was purified from total RNA using oligo(dT) magnetic beads and fragmented into 200–500 bp pieces using divalent cations at 94ºC for 8 min. The cleaved RNA fragments were copied into first-strand cDNA. After second-strand cDNA synthesis, fragments were A-tailed and indexed adapters were ligated. The products were purified and enriched by PCR to create the final cDNA library. After confirmation of successful library construction, the resulting libraries were sequenced on a HiSeq2500 or HiSEq4000 (Illumina) using a V3 flowcell generating 1 × 50 bp reads, yielding reliable results for 43, and 14 patients from groups I and ii, respectively. Optical duplicates and adapator sequences were removed from the raw sequencing reads before aligning to the transcriptome and the reference genome using TopHat 2.0 ^93^ and Bowtie 2.0 ^94^. Counts were assigned to genes using the HTSeq software package. Raw sequencing reads were mapped to the transcriptome: > 25000 different transcripts were identified that could be detected in at least 50% of the samples. For the purpose of this study, the groups of primary metastasized patients (n=14) (iii) and patients who remained disease-free for at least 6 to 10 years after the initial therapy (n=43) (i) were used for differential gene expression analysis.

#### 2D and 3D spheroid cultured cells

RNA from freshly collected cells was extracted using TRIzol™ Reagent (Thermo Scientific). RNA integrity and concentrations were measured on the Agilent Bioanalyze, and libraries were prepared using KAPA Stranded mRNA Sequencing Kit (Roche) according to the manufacturer’s instructions and as described above. After quantification with qPCR, the resulting libraries were sequenced on a HiSeq4000 (Illumina) using a flow cell generating 1×50bp single-end reads. The resulting reads were cleaned with the fastq-mcf software, after a quality control was performed with FastQC (v0.11.9). The high-quality reads were then mapped to the *Mus Musculus* reference genome (GRCm38/mm10) with HISAT2 (v2.1.0) and the abundance of reads per gene was determined with HTSeq-count. Differential gene expression analysis was performed with the R package DESeq2 (v1.22.0). To further investigate the significantly differential genes, genes were ranked according to the score-sign(log2FC)*log(padj), and used as input for a geneset enrichment analysis using GSEA software of the Broad Institute. In the pre-ranked analysis, only gene sets containing between 15 and 1000 genes were retained and the number of permutations was set at 1000. All pathways with GSEA FDR < 0.05 were considered significant. Pathway comparison and upstream regulators analysis were done in Ingenuity Pathway Analysis. The threshold criteria of P-value⍰≤ ⍰0.05 and a fold change of ≥0.5⍰in gene expression among the different pairwise comparisons were used.

### In silico gene expression analysis and pathway analysis

Transcript mRNA expression data were collected from patients with primary breast cancer from TCGA (n=1221) and METABRIC (n= 1904). Datasets were downloaded from The Cancer Genome Atlas (TCGA) portal (http://tumorsurvival.org/download.html), and cBioportal (https://www.cbioportal.org/datasets). These data were gathered from publicly available datasets and did not require institutional review board approval or patient informed consent. All patients with mRNAseq and clinical data were included. Clinical parameters, including survival information, tumor stage, tumor subtype, were acquired from TCGA’s Clinical Data Resource data set (https://gdc.cancer.gov/about-data/publications/PanCan-Clinical-2018). We used TNM staging according to the AJCC Cancer Staging Manual, 7th edition, to maintain consistency for all patients from TCGA and METABRIC datasets. Clinical information of those two datasets is shown in Supplementary Table 3. For statistical analysis, the Kaplan–Meier method was used to analyze the relationship between gene expression (CPT1A-low and CPT1A-high, median as cutoff) and survival prognosis. Univariate and multivariate Cox regressions were used to analyze the associations between patient clinical information (age, TNM stage, and tumor subtype (classified by PAM50)) and overall survival. R software (version 4.0.5 (R Project for Statistical Computing)) was used to perform KM survival analysis, univariate and multivariate Cox regressions by using a p-value < 0.05 as the filter value.

The expression levels of *CPT1A* in human tissues were analyzed in the Genevestigator database, using the HS00002-(33) public microarray and RNA-Seq study results^95^. Expression values are shown as fold change of normalized expression values calculated by Genevestigator calculated using standard normalization methods for different microarray platforms.

Pathway analysis: QIAGEN’s Ingenuity® Pathway Analysis (IPA®, QIAGEN) tool was applied to differential gene expression data obtaining from the RNA sequencing, The activation score was calculated based on the direction of change (i.e. expression in the sg*CPT1A* + acetate relative to control sh*CPT1A* without acetate. The threshold criteria of P-value⍰≤ ⍰0.05 and a fold change of ≥0.5 were used.

### Statistical analysis

Statistical data analysis was performed using GraphPad Prism 7 (GraphPad Software) on n ≥ 3 biological replicates. Details of statistical tests and post-tests are presented in the figure legends. Determination of mathematical outliers was performed using Grubb’s test. Sample size for all in vitro experiments was chosen empirically. For in vivo experiments, sample size was determined using power calculations with β = 0.8 and P < 0.05, based on preliminary data. Data are presented as mean ± s.e.m.

